# Comparative analysis of *Wolbachia* maternal transmission and localization in host ovaries

**DOI:** 10.1101/2024.03.03.583170

**Authors:** Michael T.J. Hague, Timothy B. Wheeler, Brandon S. Cooper

## Abstract

Many insects and other animals carry microbial endosymbionts that influence their reproduction and fitness. These relationships only persist if endosymbionts are reliably transmitted from one host generation to the next. *Wolbachia* are maternally transmitted endosymbionts found in most insect species, but transmission rates can vary across environments. Maternal transmission of *w*Mel *Wolbachia* depends on temperature in natural *Drosophila melanogaster* hosts and in transinfected *Aedes aegypti*, where *w*Mel is used to block pathogens that cause human disease. In *D. melanogaster*, *w*Mel transmission declines in the cold as *Wolbachia* become less abundant in host ovaries and at the posterior pole plasm (the site of germline formation) in mature oocytes. Here, we assess how temperature affects maternal transmission and underlying patterns of *Wolbachia* localization across 10 *Wolbachia* strains diverged up to 50 million years—including strains closely related to *w*Mel—and their natural *Drosophila* hosts. Many *Wolbachia* maintain high transmission rates across temperatures, despite highly variable (and sometimes low) levels of *Wolbachia* in the ovaries and at the developing germline in late-stage oocytes. Identifying strains like closely related *w*Mel-like *Wolbachia* with stable transmission across variable environmental conditions may improve the efficacy of *Wolbachia*-based biocontrol efforts as they expand into globally diverse environments.

## INTRODUCTION

Microbes form diverse relationships with host organisms that span the tree of life, including animals, plants, and protists. Endosymbiosis is an especially close relationship where microbes occupy eukaryotic host cells. Heritable endosymbionts are particularly common in insects^1–3^, and these relationships alter fundamental aspects of host biology, including reproduction^4,5^, protection for natural enemies^6–9^, and nutrient acquisition^10–12^.

Many endosymbionts rely on vertical-maternal transmission to spread and persist in insect populations^13–15^. In obligate relationships, these endosymbionts must be maternally transmitted to ensure host survival (e.g., *Buchnera* and aphids^16^). For facultative endosymbionts like *Wolbachia*, imperfect maternal transmission decreases the prevalence of *Wolbachia*-positive individuals within the host population each generation^17–21^. Thus, maternal transmission is a fundamental determinant of how endosymbionts spread, persist, and evolve in host populations^17,22,23^. In addition, endosymbiont-based programs to control human diseases^24–26^ and agricultural pests^27,28^ explicitly depend on efficient maternal transmission of *Wolbachia* in transinfected insect vectors and pest populations. This includes introducing the virus-blocking *w*Mel *Wolbachia* strain from *Drosophila melanogaster* into mosquito populations to block dengue and other arboviruses on multiple continents^29–32^. Nonetheless, it is largely unknown why maternal transmission may break down under certain circumstances, especially for non-model systems^15,21,33–36^.

Maternal transmission is predicted to depend on endosymbiont density (titer) in female reproductive tissues, and specifically at the site of germline formation during oogenesis^14^. In insects, endosymbionts often achieve vertical transmission by occupying the female ovaries. Endosymbionts are observed in germline stem cells in female hosts where they maintain a tight association with the germline throughout oogenesis, thereby ensuring a germline-to-germline route of vertical transmission through the host matriline^14,15,36,37^. In a more circuitous route, some endosymbionts may also colonize the germline from neighboring somatic cells via cell-to-cell migration through a soma-to-germline route of transmission^4,14,15,38,39^. Our canonical understanding of these cellular processes—and their contributions to maternal transmission—are based almost entirely in the context of standard laboratory conditions where maternal transmission rates are near perfect or perfect^18,20,21,40^. However, maternal transmission of facultative endosymbionts is often imperfect in natural host populations^19,20,41,42^ and the cellular processes that lead to transmission breakdown under natural conditions are generally unresolved^21,36^. Understanding why transmission breaks down under certain conditions is critical for predicting endosymbiont spread in nature and improving *Wolbachia*-based biocontrol programs^25,33,34,43^.

Maternally transmitted *Wolbachia* are the most common endosymbionts on earth, associating with about half of all insect species, as well as other arthropods and nematodes^4,44–46^. *Wolbachia* are primarily maternally (vertically) transmitted via the host germline within host species; although, host switching via horizonal transfer between host species is common on evolutionary timescales^47–54^. *Wolbachia* generally form facultative relationships (from the host perspective) in insects, with some individuals in the host population that do not carry *Wolbachia* due to imperfect maternal transmission. Maternal transmission is perhaps the most important determinant of *Wolbachia* prevalence in host populations: theory demonstrates that *Wolbachia* frequency dynamics and equilibria are approximated by the degree of imperfect maternal transmission (*μ*), the relative fitness (e.g., fecundity) of females with *Wolbachia* (*F*), and the strength of *Wolbachia*-induced cytoplasmic incompatibility (*s_h_*)^17,55^. Some *Wolbachia* strains (e.g., *w*Ri in *D. simulans*) cause cytoplasmic incompatibility (CI), a crossing incompatibility that generates a frequency-dependent advantage for females with *Wolbachia*^5,56^.

The prevalence of different *Wolbachia* strains varies widely among host systems, which may be partially explained by differences in maternal transmission rates. For instance, the strong CI-causing *w*Ri strain is observed at high frequencies in global *D. simulans* populations^17,18,57^, whereas the weak CI-causing *w*Mel strain is found at intermediate, fluctuating frequencies in *D. melanogaster*^58^. *w*Ri frequencies can be plausibly explained by strong CI that causes spread to high frequencies, whereas lower, variable *w*Mel frequencies can be plausibly explained by maternal transmission breakdown as temperatures decrease, in addition to a minimal influence of CI^21^. The thermal sensitivity of maternal transmission may be a feature of *w*Mel, given that heat stress also reduces *w*Mel transmission in transinfected *Aedes aegypti* hosts^33,34,43^. Identifying *Wolbachia* variants that are efficiently transmitted across environmental conditions could potentially contribute to improving the efficacy of *Wolbachia* based biocontrol of human diseases and agricultural pests^25,26^.

What might underlie variable maternal transmission rates? Maternal transmission breakdown is predicted to stem from reductions in *Wolbachia* abundance and localization throughout host development and specifically at the critical stages of oogenesis thought to facilitate germline-to-germline maternal transmission^14,15,21,36,59–63^. *w*Mel cells are present in the germline stem cells of adult females, and over the course of oogenesis, *w*Mel localize via kinesin-mediated transport to the pole plasm—the site of germline formation—at the posterior cortex of the developing oocyte. In late oogenesis (stage 10), *w*Mel *Wolbachia* display a pattern of posterior localization in oocytes, which is considered to be a key step in the process of germline-to-germline maternal transmission under standard laboratory conditions^14,50,59–63^. In support of this, we recently demonstrated that declining *w*Mel transmission in the cold covaries with a significant reduction in cellular *Wolbachia* abundance at the posterior cortex of stage 10 oocytes, which can plausibly explain why maternal transmission is thermally sensitive^21^.

Outside of the *w*Mel strain, it is unclear how temperature influences *Wolbachia* maternal transmission and posterior localization in stage 10 oocytes for other diverse *Wolbachia*-host associations^15,20^. Under standard 25°C conditions, divergent *Wolbachia* strains exhibit highly variable patterns of localization to the pole plasm at the posterior cortex of stage 10 oocytes of *Drosophila* species, deviating from our canonical understanding of *w*Mel^15^. These patterns range from strongly localized (e.g., *w*Ana in *D. anannassae*) to a complete lack of localization (e.g., *w*Tri in *D. triauraria*) at the site of germline formation. Some strains, particularly the divergent B-group *w*Mau strain, also exhibit evidence for an alternative soma-to-germline route of transmission, whereby *Wolbachia* in somatically derived follicle cells may invade the oocyte via cell-to-cell migration. It is entirely unknown whether the diverse localization patterns of these *Wolbachia* strains influence rates of maternal transmission and/or are sensitive to thermal perturbation like *w*Mel.

Here, we examine temperature effects on *Wolbachia* maternal transmission and localization in host tissues across 10 divergent *Wolbachia* strains naturally found in eight *Drosophila* host species within the *melanogaster* species group (Figure 1, Table S1). These *Wolbachia* comprise nine A-group *Wolbachia*—including closely related *w*Mel-like strains (*w*Mel, *w*MelCS, *w*Seg in *D. seguyi*, and *w*Cha in *D. chauvacae*) and closely related *w*Ri-like strains (*w*Ri, *w*Ana in *D. ananassae*, *w*Aura in *D. auraria*, and *w*Tri in *D. triauraria*)^20,51,53^—and divergent B-group *w*Mau in *D. mauritiana* that diverged up to 46 million years ago from the other A-group strains^40^. We reared flies at 25° and 20°C and then measured maternal transmission rates in conjunction with *Wolbachia* densities (titer) at the tissue level in female ovaries and the remaining somatic tissues. We then used confocal microscopy to examine how temperature impacts the cellular abundance of *Wolbachia* at the site of germline formation at the posterior cortex of stage 10 oocytes during oogenesis. Our results reveal diverse patterns of localization in ovary tissues and late-stage oocytes that are temperature-dependent, but only weakly predictive of maternal transmission rates. We also identify *Wolbachia* strains that are transmitted efficiently regardless of temperature.

**Figure 1.**
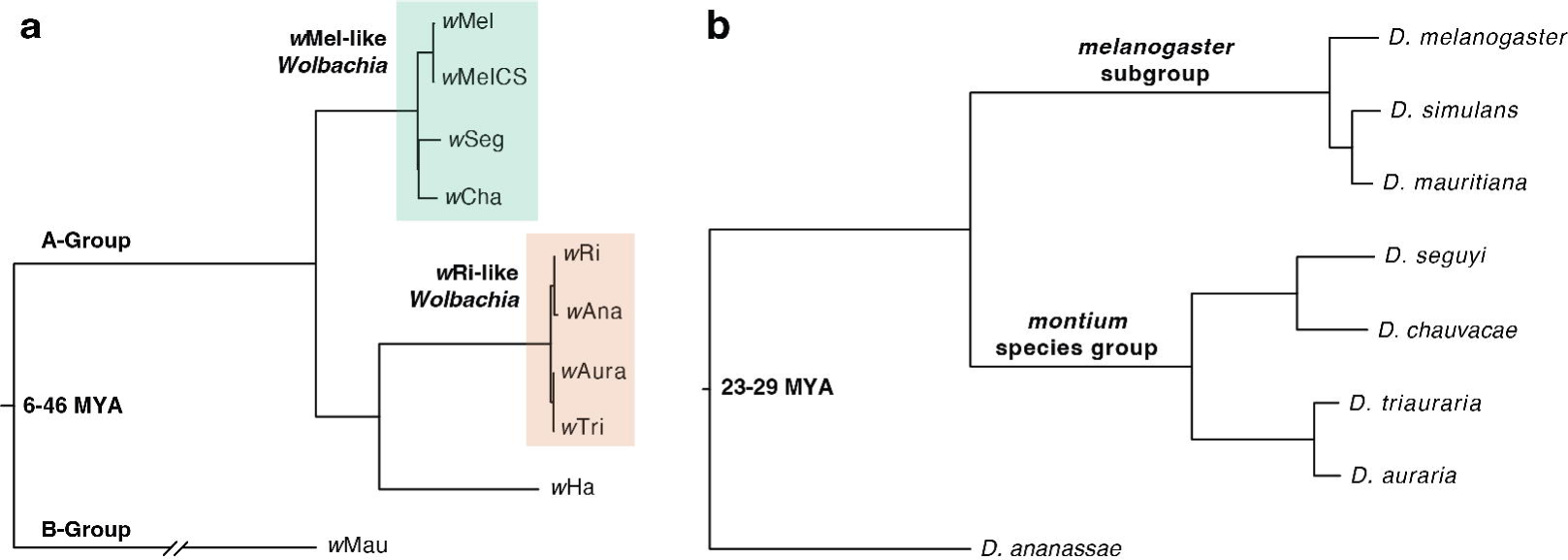
Divergent *Wolbachia* strains found in *Drosophila* host species. **(a)** Estimated Bayesian phylogram of the 10 A- and B-group *Wolbachia* strains included in the study. The phylogram was estimated using 170 single-copy genes of identical length in all genomes, spanning 135,105 bp. **(b)** Estimated Bayesian phylogram of the eight *Drosophila* species using 20 single-copy genes. All nodes on both trees are supported with Bayesian posterior probabilities of 1. Estimates of *Wolbachia* and *Drosophila* divergence reported in millions of years ago (MYA) are reproduced from Meany *et al.*^40^ and Suvorov *et al.*^136^, respectively. The *Wolbachia* and host trees are generally discordant, as expected with frequent *Wolbachia* host switching^47,48,51,53,54^.

## RESULTS

We estimated the mean rate of maternal transmission (± BC_a_ confidence intervals) for each *Wolbachia*-host genotype at 25° and 20°C by pairing individual virgin females with *Wolbachia*-free males (see Methods), allowing the females to lay for eight days, and then measuring maternal transmission (1 – *μ*) to the newly emerged offspring of each subline (Figure 2, Table S2)^20,21^. The final dataset comprised 365 F0 females and 3,612 F1 offspring. Overall, we found a significant interaction effect between the *Wolbachia*-host genotype and temperature (GLMM LRT, χ^2^_(9)_ = 23.49, *P* = 0.005), such that temperature effects on maternal transmission varied depending on the *Wolbachia*-host genotype. For most *Wolbachia*-host genotypes, mean rates of maternal transmission were perfect (e.g., *w*Ri-*D. simulans*) or near-perfect (e.g., *w*Mel-like *w*Seg-*D. seguyi*) and invariant across the two temperatures (Figure 2, Table S3), with the exception of a few specific *Wolbachia*-host genotypes. Maternal transmission of *w*Mel by *D. melanogaster* (*W* = 251.5, *P* = 0.003) and *w*Aura by *D. auraria* (*W* = 154, *P* = 0.038) decreased significantly at 20°C; although, *w*Mel experienced a much greater decline in the average rate of transmission (14.9%) than *w*Aura (2.8%). In contrast, B-group *w*Mau transmission by *D. mauritiana* increased slightly by 3.4% in the cold at 20°C (*W* = 117, *P* = 0.019). Notably, our analysis also revealed a significant effect of sex (GLMM LRT, χ^2^_(1)_ = 63.71, *P* < 0.001), whereby rates of maternal transmission tended to be lower to female offspring than males (Table S2). This finding is consistent with our previous findings for *w*Mel^21^ and the *w*Mel-like *w*Yak strain in *D. yakuba*^20^, but counter to theoretical expectations that selection should favor faithful transmission to female offspring^64,65^.

**Figure 2.**
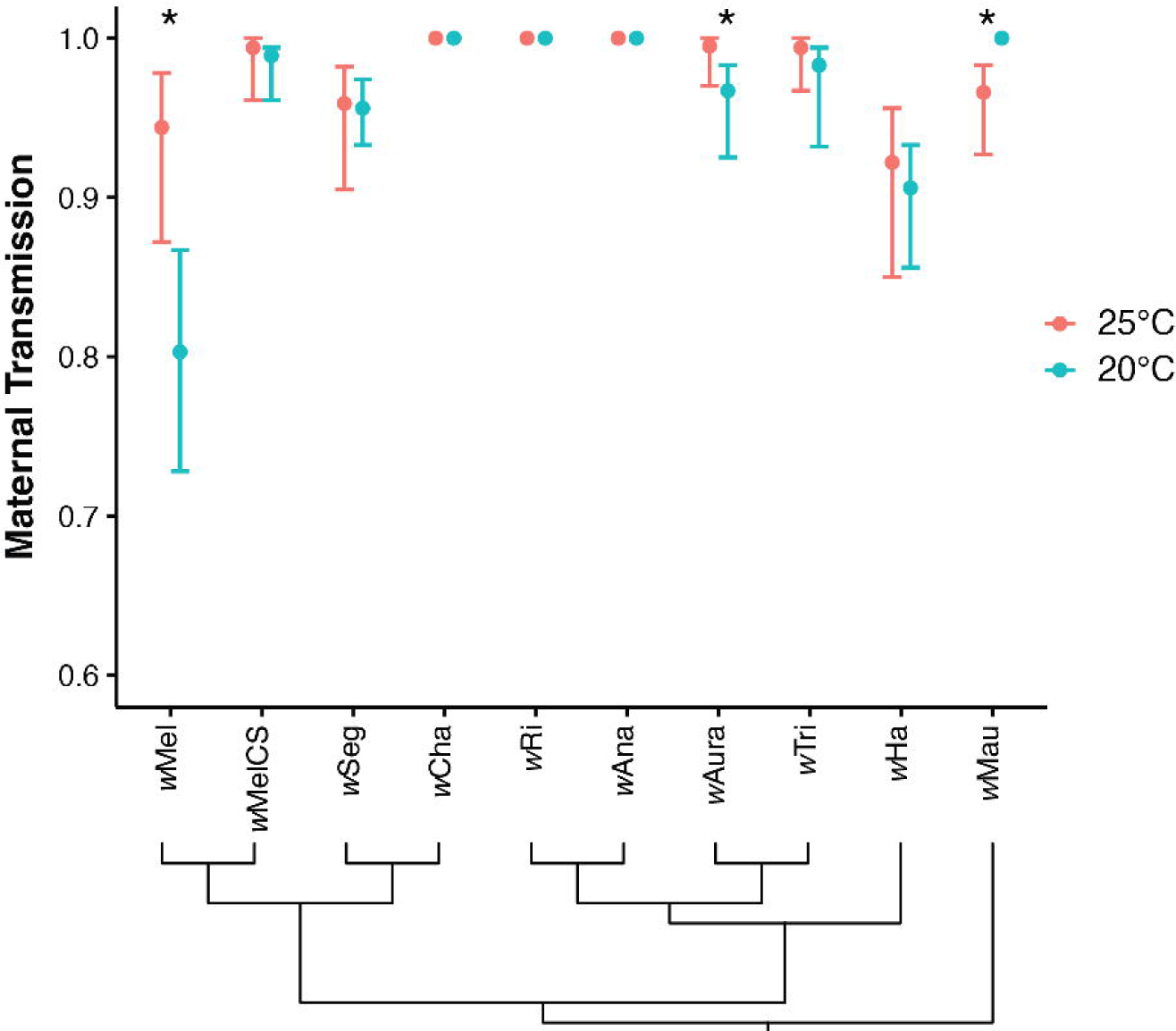
Maternal transmission is high and stable across temperatures for many *Wolbachia* strains. Mean maternal transmission rates (±BC_a_ confidence intervals) of different *Wolbachia* strains in their naturally associated host species (*N* = 364 sublines). Asterisks indicate the rates of maternal transmission differ between 25° and 20°C for a given *Wolbachia* strain according to a Wilcoxon rank-sum test at *P* < 0.05. Below, the cladogram depicts evolutionary relationships among *Wolbachia* strains.

To test for temperature effects on *Wolbachia* at the tissue level, we dissected out the ovaries from a subset of the same F0 females assessed for maternal transmission above (*N* = 183). For this analysis we used qPCR to measure *Wolbachia* densities in the ovaries and the remaining somatic carcass tissue (Figure 3). We found a significant interaction effect between the *Wolbachia*-host genotype and temperature on *Wolbachia* density in ovaries (Two-way ANOVA; *F*_(9,161)_ = 4.286, *P* < 0.001) and the remaining carcasses (*F*_(9,161)_ = 10.612, *P* < 0.001), such that temperature effects on *Wolbachia* density varied depending on the *Wolbachia*-host genotype. While we uncovered considerable variation depending on the *Wolbachia*-host genotype and temperature, *Wolbachia* ovary densities generally tended to decline at 20°C for the majority of systems (Figure 3, Table S4). Five of the *Wolbachia* strains (*w*Mel, *w*Ri, *w*Ana, *w*Ha, and *w*Mau) decreased in density in the ovaries in the cold. In contrast, carcasses exhibited variation in the directionality of changes in *Wolbachia* density. *w*Mel, *w*MelCS, *w*Seg, and *w*Aura densities decreased significantly in the cold, whereas *w*Cha and *w*Mau significantly increased. *w*Cha densities in carcasses were roughly an order of magnitude lower than all other strains, regardless of temperature, suggesting a relatively high degree of localization to the ovaries. *w*Mel, the strain that experienced the most dramatic decline in transmission in the cold (Figure 2), was also the only *Wolbachia* strain that experienced a significant decline in *Wolbachia* density in both the ovaries and the carcasses in the cold. This included a particularly large decline in density in the carcasses (a 4.7-fold reduction) compared to the other strains.

**Figure 3.**
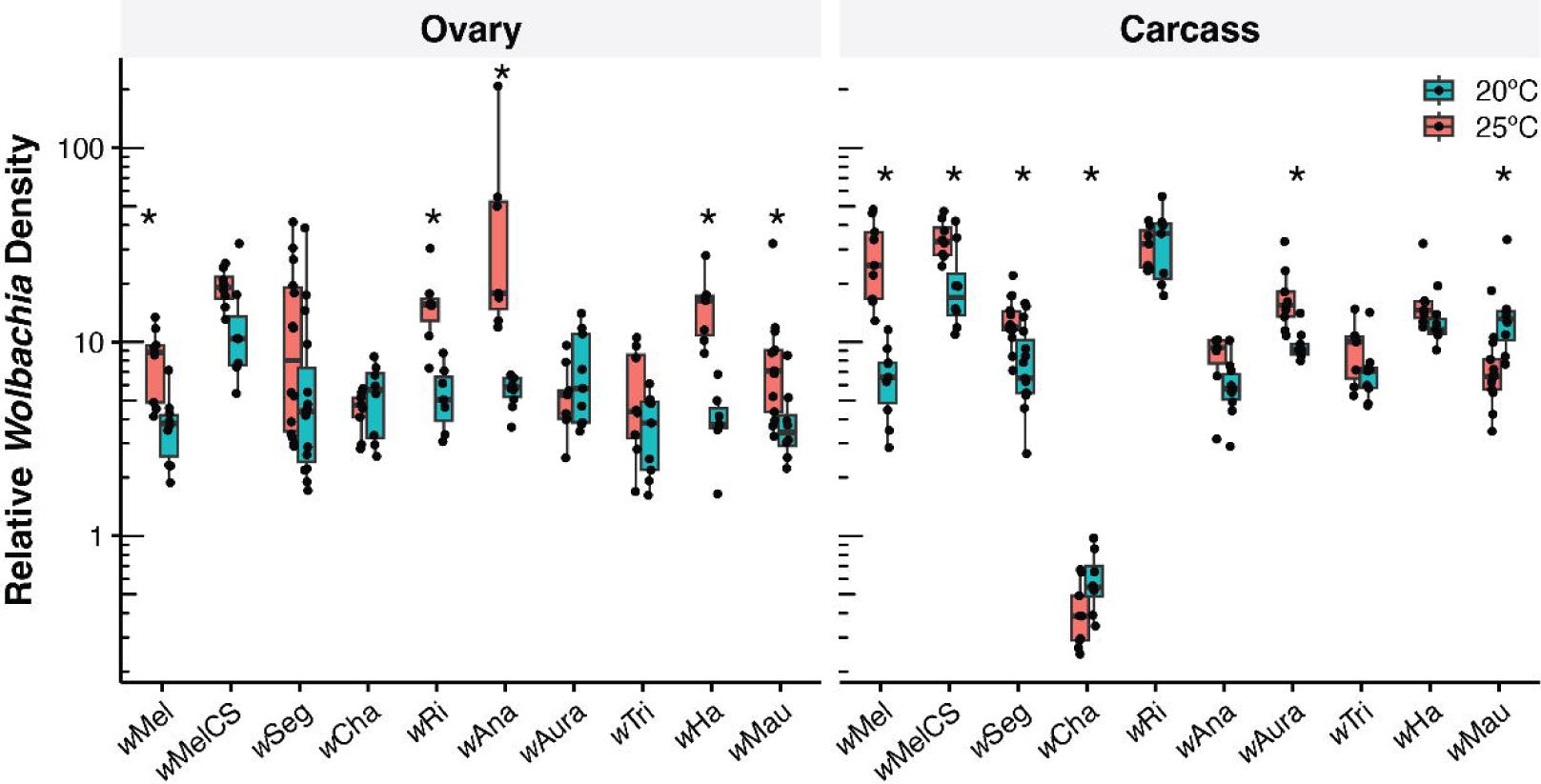
*Wolbachia* densities in host tissues vary by *Wolbachia*-host system and temperature. *Wolbachia* densities are shown for dissected female ovaries and the remaining somatic carcass tissue (*N* = 183 F0 females). Asterisks indicate the *Wolbachia* density differs between 25° and 20°C for a given *Wolbachia* strain according to a Wilcoxon rank-sum test at *P* < 0.05. Boxplots show medians, first and third quartiles (hinges), and the smallest/largest values within 1.5*IQR of the hinges (whiskers).

For the three *Wolbachia*-host systems with temperature-dependent transmission (*w*Mel, *w*Aura, and *w*Mau), we also tested for a correlation between transmission rates and *Wolbachia* density in the ovaries and carcasses of the same individual females (Figure S1). Declining *w*Mel transmission in the cold was significantly correlated with *Wolbachia* density in the ovaries (Spearman’s *ρ* = 0.537, *P* = 0.022), but not the carcasses (*ρ* = 0.392, *P* = 0.108), providing additional support that reduced *w*Mel density in reproductive tissues contributes to transmission breakdown in the cold. Maternal transmission of *w*Aura by *D. auraria* did not strongly correlate with *w*Aura density in the ovaries (*ρ* = -0.244, *P* = 0.380) or the carcasses (*ρ* = 0.035, *P* = 0.902). Finally, increasing maternal transmission of *w*Mau by *D. mauritiana* in the cold did not correlate with *w*Mau density in the ovaries (*ρ* = -0.119, *P* = 0.649), but was significantly correlated with density in the carcasses (*ρ* = 0.549, *P* = 0.022).

Finally, we examined how temperature impacts cellular *Wolbachia* abundance at the site of germline formation in stage 10 oocytes during host oogenesis. At this stage, *w*Mel *Wolbachia* localization at the posterior cortex of the oocyte is considered to be a critical step in the process of germline-to-germline *Wolbachia* maternal transmission^14,15,21,36,59–63^. We imaged a total of 283 stage 10 oocytes from eight-day-old females reared at the two different temperatures and quantified cellular *Wolbachia* abundance (measured as corrected total cell fluorescence; CTCF) in the whole oocytes, the posterior region, and the posterior cortex (Figure S2)^14,15,21^. We defined the posterior region as the posterior 1/8^th^ portion of the oocyte and the posterior cortex as the narrow cortical region of Vasa expression (a germline protein component)^15,63^. For the sake of clarity and consistency with the literature, we refer to our *Wolbachia* quantifications with confocal microscopy here as “cellular *Wolbachia* abundance” in oocytes, as opposed to the quantification of “*Wolbachia* density” at the tissue level using qPCR (described above).

Cellular *Wolbachia* abundance in whole oocytes (Two-way ANOVA; *F*_(9,263)_ = 2.9, *P* = 0.003), at the posterior region (*F*_(9,263)_ = 4.16, *P* < 0.001), and at the posterior cortex (*F*_(9,263)_ = 2.41, *P* = 0.012) depended on significant two-way interactions between the *Wolbachia*-host genotype and temperature. As with *Wolbachia* density, cellular *Wolbachia* abundance in oocytes varied considerably depending on the *Wolbachia*-host genotype and temperature (Figures 4, S3). The cellular abundance of six of the *Wolbachia* strains (*w*MelCS, *w*Seg, *w*Ana, *w*Aura, and *w*Tri, and *w*Ha) declined significantly throughout whole oocytes in the cold (Table S5). Temperature affected fewer strains (only *w*Mel, *w*MelCS, *w*Ri, and *w*Aura) at the posterior region and the posterior cortex, the location of the developing germline. *w*Mel and *w*Aura, the two strains transmitted at significantly lower rates in the cold, both declined significantly in abundance in the posterior oocyte region (*W* = 113, *P* = 0.015; *W* = 172, *P* = 0.041). *w*Mel cellular abundance also significantly declined at the posterior cortex (*W* = 122, *P* = 0.002). Interestingly, *w*MelCS also exhibited a significant decline in cellular abundance in the posterior region (*W* = 203, *P* < 0.001) and the cortex (*W* = 205, *P* < 0.001), despite the fact that *w*MelCS transmission by *D. melanogaster* was not altered in the cold (Figure 2). This can perhaps be explained by the fact that, of all 10 strains, *w*MelCS had the highest cellular abundance at the posterior region and cortex, regardless of temperature. Finally, *w*Ri was perfectly transmitted by *D. simulans* at both temperatures, but *w*Ri abundance increased at the oocyte posterior region (*W* = 45, *P* = 0.004) and the cortex (*W* = 49, *P* = 0.008) in the cold. Also of note, *w*Aura and *w*Tri occurred at particularly low abundances in the oocyte posterior region and the posterior cortex, regardless of temperature, despite both being transmitted at relatively high rates by their female hosts (1 – *μ* > 0.967).

**Figure 4.**
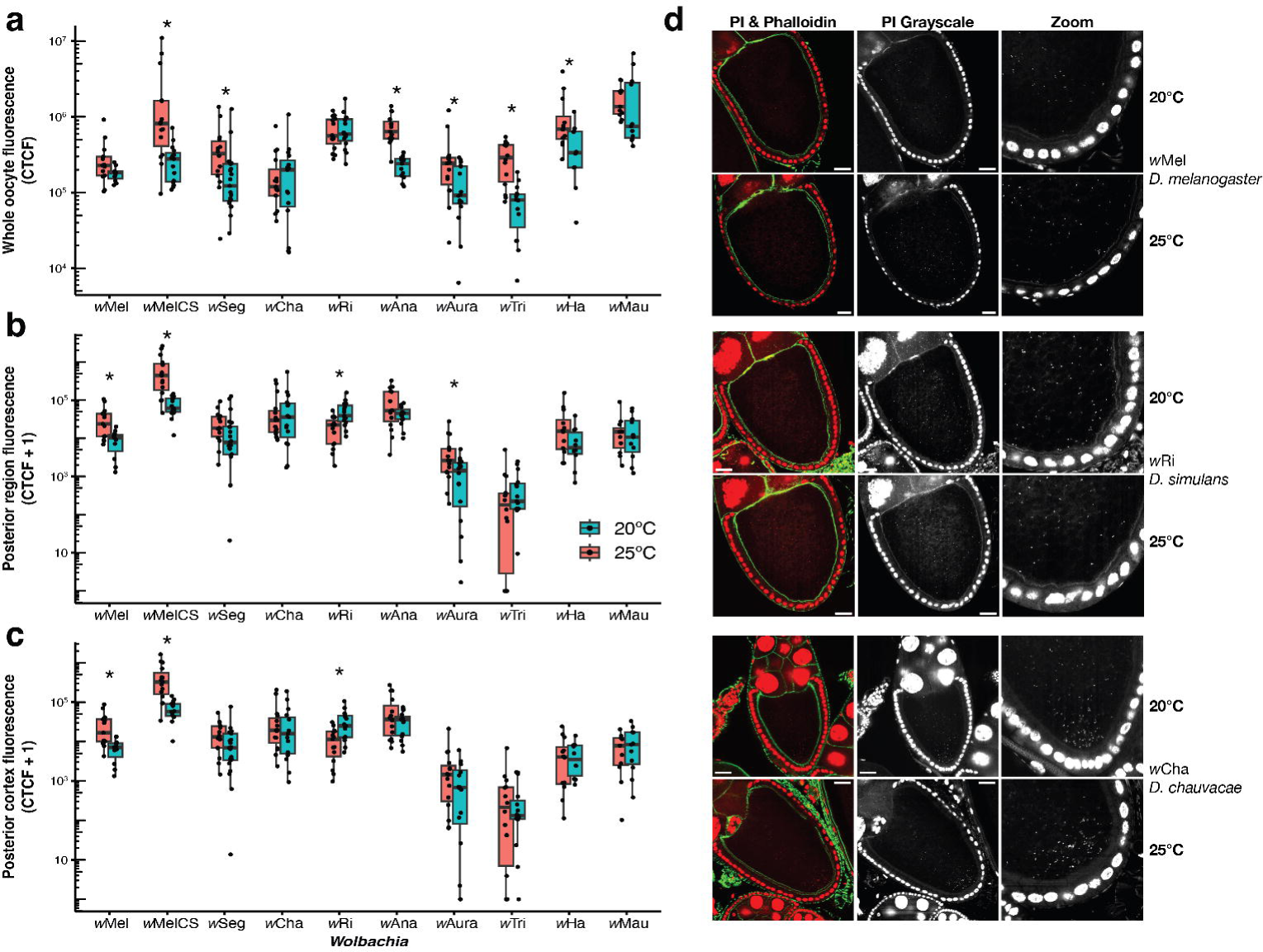
Cellular *Wolbachia* abundance in late-stage oocytes generally does not align with maternal transmission rates. Cellular *Wolbachia* abundance in stage 10 oocytes (measured as fluorescence due to propidium iodide; CTCF), measured in **(a)** the whole oocyte, **(b)** the posterior region, and **(c)** the posterior cortex (*N* = 283 oocytes). Asterisks indicate that *Wolbachia* abundance differs between 25° and 20°C for a given *Wolbachia* strain according to a Wilcoxon rank-sum test at *P* < 0.05. **(d)** Representative confocal images of *Wolbachia* strains that have decreased abundance at the posterior cortex in the cold (*w*Mel), increased abundance at the cortex in the cold (*w*Ri), and no change in the cold (*w*Cha). Confocal micrographs are DNA-stained with PI (red) and actin-stained with phalloidin (green). The second column depicts a single channel image of the PI stain and the third column depicts an enlarged PI-stained image at the posterior cortex of each oocyte. Scale bars are set to 25 µm.

Our analyses revealed that the diverse *Wolbachia* strains and *Drosophila* host species exhibit extensive variation in *Wolbachia* density in host tissues (Figure 3) and abundance in host oocytes (Figure 4). We hypothesized that this diversity may be explained by factors in the *Wolbachia* or host genomes (or both). If so, we expect closely related *Wolbachia* strains (or host species) to exhibit similar patterns of *Wolbachia* localization in host tissues^15,66,67^. We used the *Wolbachia* and host trees (Figure 1) to test whether patterns of *Wolbachia* localization in tissues and oocytes exhibit phylogenetic signal using Pagel’s λ^68^. A value of 1 is consistent with trait evolution that entirely agrees with the *Wolbachia* (or host) phylogeny, suggesting factors in the *Wolbachia* genome (or host) contribute to *Wolbachia* localization patterns (i.e., strong phylogenetic signal). In contrast, a value of 0 is consistent with trait evolution that occurs independently of phylogenetic relationships^68,69^. We found no evidence that *Wolbachia* density in the ovaries and carcasses, nor cellular abundance in the oocytes, exhibit phylogenetic signal on the *Wolbachia* phylogeny (Table S6, Figure S4), implying that factors in the *Wolbachia* genome do not account for observed diversity. In contrast, we found that *Wolbachia* density in the ovaries at 25°C exhibits strong, significant phylogenetic signal on the host phylogeny (λ = 1.000, *P* = 0.001), implying that factors in the host genome may help determine observed diversity of *Wolbachia* densities in host ovaries. In this instance, related species in the *montium* species group tended to have lower *Wolbachia* densities in the ovaries relative to the *melanogaster* subgroup (see Figure S5). Divergent *D. ananassae* also had a higher *Wolbachia* density than all the other host species. We also found that cellular *Wolbachia* abundance in the oocyte posterior region (λ = 1.000, *P* = 0.037) and the posterior cortex (λ = 1.000, *P* = 0.036) exhibited a significant signature of phylogenetic signal at 20°C on the host phylogeny. This pattern is largely driven by the relatively low abundance of *Wolbachia* in the posterior oocytes of the two closely related species *D. triauraria* and *auraria*.

## DISCUSSION

Maternal transmission rates are a key determinant of *Wolbachia* prevalence in host populations^17,55^. For many *Wolbachia* strains and host species, we found that maternal transmission is generally high and stable across two ecologically relevant temperatures, 25° and 20°C, that flies are likely to experience in natural environments^21,70^. Two *Wolbachia* strains (*w*Aura and B-group *w*Mau) exhibited minor temperature-dependent changes in transmission, but *w*Mel stood out with a relatively large decrease in transmission as temperature dropped from 25° to 20°C (Figure 2). Maternal transmission is generally expected to depend on *Wolbachia* density in host reproductive tissue^20,34,43,71^, and specifically within developing oocytes^14,15,21,36,59–63^. Our analyses uncovered considerable, significant variation in *Wolbachia* densities at the tissue level and cellular abundances in oocytes, which also often varied depending on temperature. With the exception of *w*Mel, *Wolbachia* densities in ovary tissue generally did not predict maternal transmission rates (Figures 3, S1). In stage 10 oocytes, declines in the cellular abundance of *w*Mel and *w*Aura at the posterior region coincided with declining transmission in the cold (Figure 4), but the *w*MelCS and *w*Ri strains also experienced changes in cellular abundance that did not correlate with differences in maternal transmission. Together, these results suggest that maternal transmission rates cannot be accurately predicted from measurements of *Wolbachia* quantities in ovary tissue or late-stage oocytes. First, we explore the diverse patterns of *Wolbachia* localization in relation to maternal transmission in our comparative analyses. Then, we examine why maternal transmission of some strains, particularly *w*Mel, may be impacted by temperature.

At the tissue level, we found that *Wolbachia* densities in the ovaries generally did not predict rates of maternal transmission, with the exception of *w*Mel. *w*Mel density in the ovaries of individual females was tightly correlated with declining transmission rates in the cold (*ρ* = 0.537, Figure S1). Temperature did not alter *w*Aura density in the ovaries despite declining *w*Aura transmission in the cold, and *w*Mau density counterintuitively declined at 20°C despite transmission increasing. Other strains (e.g., *w*Ri and *w*Ha in *D. simulans*) experienced large declines in *Wolbachia* ovary density at 20°C with no impact on maternal transmission. *Wolbachia* densities in the remaining somatic carcass tissue also generally did not predict declining transmission in the cold; however, *w*Mau densities in the carcasses of individual females were strongly correlated with increased transmission in the cold (*ρ* = 0.549; discussed further below). Notably, *w*Cha was exceptional in terms of its very low density in carcass tissue relative to other strains, despite perfect transmission at both temperatures. Changes to host ploidy in the ovaries could plausibly contribute to the observed differences in relative *Wolbachia* density^72^; however, we did not find evidence for significant changes in host gene copy number in the ovaries across temperature treatments (Figure S6).

At the cellular level, *Wolbachia* abundance in stage 10 oocytes may help explain why *w*Mel and *w*Aura experienced declining transmission in the cold. Many strains experienced declines in cellular *Wolbachia* abundance throughout the whole oocyte at 20°C (Figure 4a), but this was generally unrelated to transmission rates. *w*Mel was one of the few strains that had a significant reduction in cellular abundance at the posterior region (Figure 4b) and cortex (Figure 4c), the site of germline formation. Similarly, *w*Aura had a reduced abundance in the posterior region at 20°C (but not at the cortex). *w*MelCS (closely related to *w*Mel) was the only additional strain with significant reductions in abundance at the posterior region and the cortex. However, *w*MelCS occurred at a much higher absolute abundance than *w*Mel, *w*Aura, and all other *Wolbachia* strains, which may explain why transmission remained high at 20°C (discussed further below). We conjecture that there is presumably a minimum cellular abundance required for efficient transmission, and *Wolbachia* strains like *w*MelCS that are generally very abundant are less likely to fall below this threshold. A reduction in cellular abundance at the oocyte posterior implies that fewer *Wolbachia* cells will be incorporated into the germline later during host development, perturbing germline-to-germline maternal transmission^21^. The fact that both *w*Mel and *w*Aura had significantly reduced abundances in the posterior region at 20°C implies that perturbations of cellular *Wolbachia* abundance in this region may help explain why some *Wolbachia* strains have reduced transmission in the cold. *w*Ri was the only other *Wolbachia* strain with an altered abundance at the oocyte posterior region and cortex, but cellular abundance increased at 20°C. This may help explain why maternal transmission of *w*Ri remained perfect at 20°C. Interestingly, a low absolute abundance of *Wolbachia* was not necessarily associated with declining maternal transmission. The *w*Tri strain (closely related to *w*Aura) occurred at a lower posterior abundance than all other strains, despite maintaining a high rate of transmission at both temperatures.

It is important to note that our analysis of *Wolbachia* in ovary tissues and stage 10 oocytes of eight-day-old females represents only a snapshot of host development. Many other unexplored factors may contribute to transmission rate variation. Other stages of development, like embryogenesis, could also be involved in mediating maternal transmission. For instance, temperature could impact *Wolbachia* localization at the newly formed germline of late blastoderm and cellularized embryos, which could influence maternal transmission^15,60^. Cellular *Wolbachia* abundance changes over the course of oogenesis and embryogenesis^15,61,72,73^ and a number of studies report complex, age-related changes in *Wolbachia* densities over the course of the host’s lifespan^74–81^. Maternal age and the order of egg-laying could also conceivably influence *Wolbachia* transmission and densities in offspring^82^. Our work motivates future studies on how temperature affects the distribution of *Wolbachia* in host tissues over the course of host development in relation to maternal transmission.

Our comparative analysis highlights how *w*Mel seems to be especially susceptible to cooling temperatures^21^, which is consistent with our previous work suggesting the strain is less prevalent in temperate host populations due to declining transmission in the cold^21,58^. In transinfected mosquitoes, *w*Mel maternal transmission and densities are also more susceptible to heat stress relative to another *Wolbachia* strain, *w*AlbB^25,26,33,34,43^. A number of factors could plausibly help explain why *w*Mel maternal transmission is especially impacted by the cool temperature, relative to other strains. At the tissue level, *w*Mel was the only *Wolbachia* we examined that exhibited a significant decline in *Wolbachia* density in both the ovaries and the carcasses at 20°C (Figure 3). Declining transmission was highly correlated with declining *w*Mel density in the ovaries, plus *w*Mel stood out with a dramatic decline in *Wolbachia* density in carcasses at 20°C compared to other strains. While *w*Mel is generally thought to follow a germline-to-germline route of maternal transmission during oogenesis, our recent work and others suggests that the strain may also rely on a soma-to germline mode of transmission^15,83,84^. *w*Mel *Wolbachia* are capable of cell-to-cell migration^84^ and *w*Mel injected directly into the abdomen of adult *D. melanogaster* females can migrate to and occupy the germline and follicle stem cells^83^. The declines of *w*Mel densities throughout host tissues could potentially impact routes of transmission originating in somatic tissue. At the cellular level, *w*Mel is also one of the few strains (in addition to *w*Aura) to experience a decline in abundance at the posterior of stage 10 oocytes, which is expected to impact a germline-to-germline route of transmission. The concurrent reduction in *Wolbachia* density in female tissues and oocytes may be a one-two punch that impacts both soma- and germline-to-germline routes of *w*Mel transmission.

The juxtaposition between *w*Mel and *w*MelCS is interesting, because the two strains diverged recently in only the last 3,000-14,000 years^85^. The *w*Mel and *w*MelCS strains we used in our study are highly similar (0.007% third-position pairwise differences). Despite the close relationship, *w*MelCS transmission was not impacted by temperature and the strain occurred at a higher cellular abundance at the oocyte posterior than any of the other strains that we examined, regardless of temperature. Some oocytes contained especially large quantities of *w*MelCS throughout the whole oocyte and at the posterior (Figure S7), as compared to all the other oocytes we examined (e.g., Figure S3). *w*MelCS is only found at low frequencies in a few populations of *D. melanogaster*^75,85^, which is attributed to the fact that ancestral *w*MelCS was largely replaced by *w*Mel in global populations of *D. melanogaster* in roughly the last 50 to 100 years^86–89^. Perhaps deleterious host effects associated with a high cellular *w*MelCS abundance in oocytes may have contributed to the global replacement of *w*MelCS by *w*Mel. To ensure maternal transmission, *Wolbachia* cells must maintain an association with the host germline without perturbing highly conserved processes of germline and oocyte development^15,62,63,73^. Migration of *w*Mel to the posterior germ plasm is coincident with recruitment of host factors required from germline formation and anterior/posterior and dorsal/ventral axis formation^62^ and an excessively high *Wolbachia* abundance can disrupt dorsal/ventral axis determination^60,62,73,90^. Perhaps similar deleterious effects may help explain why the highly abundant *w*MelCS strain was replaced by *w*Mel, despite a higher rate of transmission in the cold. Previous works has shown that higher densities of *w*MelCS in *D. melanogaster* often covaries with a reduced lifespan^75^. The cellular abundance and fitness consequences of *w*MelCS (as compared to *w*Mel) may also be influenced by interactions with the host genome and the environment^20,21,91^. Indeed, our phylogenetic analyses suggest that the host genome contributes to *Wolbachia* abundance in oocytes (Table S6). Given that interactions with *Wolbachia* and the environment influence host fitness (e.g., fecundity and longevity), it is difficult to point to a single explanation for why *w*MelCS was globally replaced by *w*Mel.

The divergent B-group *w*Mau strain was the only strain where transmission significantly increased at 20°C (Figure 2). Transmission was high at 25°C and increased to perfect at 20°C. Our previous work suggests the *w*Mau strain may also be an outlier in its mode of transmission during oogenesis relative to A-group *Wolbachia*, which tend to exhibit evidence of a germline-to-germline route of transmission^15^. *w*Mau occurs at relatively low abundance at the posterior cortex of stage 10 oocytes (Figure 4), but *w*Mau cells are frequently found within the somatically derived follicle cells surrounding the oocytes (Figure S7)^15^, which is consistent with a soma-to-germline route of transmission. The presence of *Wolbachia* in the follicle cells is generally not the case for A-group *Wolbachia* (with the exception of *w*Mel, discussed above). We found that *w*Mau was also one of the few strains (in addition to *w*Cha) where *Wolbachia* density in the somatic carcass tissues significantly increased at 20°C (Figure 3), which may help explain why transmission increased slightly at 20°C. *w*Mau transmission was also strongly correlated with *Wolbachia* densities in the carcasses of individual females. Notably, our previous analysis of two other *w*Mau-*D. mauritiana* genotypes using three- to five-day old females found perfect rates of transmission at 25°C^40^, unlike our results here (Figure 2). This suggests that differences in fly age^81^, the *Wolbachia* and host genomes^21^, or other unknown factors may also contribute to variation in rates of maternal transmission.

The *w*Mau findings also may inform our previous work on the thermoregulatory behavior of *Drosophila* host species. We found that *Drosophila* species carrying A-group *Wolbachia*, including *w*Ri and *w*Ha, tend to prefer cooler temperatures on a thermogradient (relative to flies without *Wolbachia*), whereas *D. mauritiana* flies carrying *w*Mau prefer warmer temperatures^66^. Taken together, these results suggest that fly movement to cool temperatures reduces the density of A-group *Wolbachia* in host tissues, whereas movement to warm temperatures reduces B-group *w*Mau density. This is consistent with the inference that flies may thermoregulate as a behavioral response to ameliorate negative effects of infection and a high *Wolbachia* density^67,92–95^. Movement to cooler temperatures with A-group *Wolbachia* (i.e., behavioral chill) and to warmer temperatures with B-group *w*Mau (behavioral fever) are both behaviors that would reduce *Wolbachia* density in host tissues according to our results here (Figure 3). This finding motives further work dissecting the relationships among thermoregulatory behavior, *Wolbachia* density in host tissues, and maternal transmission.

Our phylogenomic analyses revealed that *Wolbachia* density in the ovaries at 25°C and cellular *Wolbachia* abundance in the oocyte posterior region and cortex at 20°C all exhibit phylogenetic signal on the host phylogeny (Table S6, Figure S5), implying that factors in the host genome help explain the observed diversity in *Wolbachia* localization across species. For example, interspecific variation in the host proteins that *Wolbachia* engage during oogenesis could plausibly influence *Wolbachia* localization patterns at the posterior cortex in stage 10 oocytes^15^. Unfortunately, it is not yet known how *Wolbachia* engage host factors like kinesin during oogenesis^63^. Other work also suggests the host genome can influence *Wolbachia* abundance in oocytes^21^ and embryos^60^. Similarly, genome-wide screens reveal that host factors play an important role in determining cellular *Wolbachia* abundance^96,97^. The findings here complement our previous comparative analyses that suggest the *Wolbachia* genome also contributes to variation in *Wolbachia* localization patterns in oocytes^15^ and effects on host thermoregulatory behavior^66^. While we did not identify any traits that exhibit phylogenetic signal on the *Wolbachia* phylogeny, it is worth noting that relatively few *Wolbachia* genomic changes (e.g., in the ampliconic “Octomom” gene region) are required to influence density and tissue distributions^75,98–101^. This highlights the potential for rapid changes in these traits and their effects on transmission. DNA analyses reveal discordant *Wolbachia* and host phylogenies, with recently diverged *Wolbachia* found in distantly related hosts^47–54,102^. This includes evidence of the rapid spread of *w*Ri-like *Wolbachia* across *Drosophila* flies diverged about 50 MYA^51^ and *w*Mel-like *Wolbachia* across holometabolous insects^103^ diverged about 350 MYA^53^. Given the short persistence of *Wolbachia* with hosts in our study (see Figure S8), it is unlikely that the observed patterns of *Wolbachia* transmission and tissue localization result from coevolution between *Wolbachia* and host genomes.

The rate of *Wolbachia* maternal transmission, in conjunction with *Wolbachia* effects on host fitness and cytoplasmic compatibility (CI), ultimately determine *Wolbachia* prevalence in host populations^17,55^. In this study, we uncovered limited variation in maternal transmission rates across temperatures, despite dramatic variation in *Wolbachia* localization patterns within host tissues. Nonetheless, a few *Wolbachia* strains, particularly *w*Mel, had variable transmission rates that are likely to impact *Wolbachia* prevalence in natural host populations. For instance *w*Mel tends to cause weak CI^81,104,105^ and occurs at lower frequencies in temperate host populations in Australia and North America, which can plausibly be explained by declining maternal transmission at cool temperatures (Figure 2)^21^. Similarly, *w*Mau does not cause CI and occurs at intermediate frequencies in *D. mauritiana* on Mauritius^40,106^. Our results suggest temperature-related variation in *w*Mau transmission may perhaps help contribute to intermediate frequencies. Supporting our prior observations^20,21^, *Wolbachia* transmission rates to female offspring were lower than to male offspring (Table S2), which could influence sex-specific *Wolbachia* frequencies observed in nature. Future work focused on understanding the causes of sex-specific transmission rates and *Wolbachia* densities will be important for understanding this pattern.

Other *Wolbachia* strains like *w*Ri^17–19,107^, *w*Ha^108–110^, and *w*Ana^111,112^ cause strong CI and tend to occur at high equilibrium frequencies in nature. In addition to strong CI, our results suggest that stable transmission rates may help these strains maintain high frequencies in the face of fluctuating conditions (Figure 2). However, this may not always be the case, as we found that maternal transmission the strong CI-causing strain *w*Aura^51,111^ can be perturbed by temperature. Unlike CI and *Wolbachia* maternal transmission, the role of *Wolbachia* effects on host fitness in natural *Drosophila* populations is poorly understood, although recent work suggests *Wolbachia* block viruses in their native *Drosophila* hosts^8,9,113,114^.

Understanding how temperature influences *Wolbachia* maternal transmission, especially in relation to other key factors like CI and host fitness effects, is critical for explaining global *Wolbachia* prevalence in host populations. Research on this front will also inform *Wolbachia*-based biocontrol programs, as thermally stable *Wolbachia* strains can be leveraged to improve the efficacy of *Wolbachia*-based biocontrol applications in mosquito populations that experience extreme environments^25,115^. In this respect, temperature and the environment are emerging as key factors that mediate interactions between *Wolbachia*, their hosts, and pathogens.

## METHODS

### Fly lines

We evaluated 10 different *Wolbachia* strains that naturally occur in eight different *Drosophila* species (Table S1). For two of these host species, we tested multiple *Wolbachia*-host genotypes: *w*Mel and *w*MelCS in *D. melanogaster* and *w*Ri and *w*Ha in *D. simulans*. With the exception of the *w*MelCS-*D. melanogaster* line, all the *Wolbachia*-host genotypes were sampled from nature to form isofemale lines, such that single gravid females were collected from the field and placed individually into vials (see Hague et al., 2020^66^ for further discussion of the *w*MelCS genotype). Each isofemale line was stably maintained in the lab for at least four years prior to the experiments. For the maternal transmission experiments explained below, we paired each *Wolbachia*-positive female with *Wolbachia*-free males of the same host species (Table S1). Here, we used males from naturally *Wolbachia*-free isofemale lines whenever possible; however, in some cases, we were unable to obtain natural genotypes without *Wolbachia*. For these species, *Wolbachia*-free genotypes were created by treating the *Wolbachia*-positive genotype with 0.03% tetracycline for at least four generations. After the fourth generation, we used quantitative PCR (qPCR) to confirm that flies were cleared of *Wolbachia*^66^. We then reconstituted the gut microbiome of the tetracycline-cleared flies by rearing them on food where *Wolbachia*-positive males of the same genotype had been fed and defecated for the prior 48 hours. Tetracycline-cleared flies were given at least three more generations before we conducted experiments to avoid detrimental effect of the antibiotic treatment on mitochondrial function^116^.

### Maternal transmission

We tested how temperature influences maternal transmission and *Wolbachia* distributions in host tissues by rearing females of each genotype at a moderate (25°C) and low (20°C) temperatures. Prior to experiments, the isofemale lines were reared for at least two generations in incubators set to each temperature and a 12L:12D light cycle (Percival Model I-36LL) on a standard food diet^66^. We estimated maternal transmission at each temperature by aspirating individual virgin females into vials with two 3-5 day old virgin males of the *Wolbachia*-free genotype and then allowing females to lay eggs for eight days^20,21^. We crossed *Wolbachia*-positive females to males without *Wolbachia*, because in crosses with *Wolbachia*-positive males, *Wolbachia*-free ova produced by *Wolbachia*-positive females could be lost if they are susceptible to cytoplasmic incompatibility, potentially leading to an overestimation of maternal transmission^18^. The males in our experiments were all collected from the *Wolbachia*-free genotypes that were maintained at 25°C.

Newly emerged adult F1 offspring were collected in the first 48 hours of emergence, preserved in 95% ethanol, and stored at -20°C for later analysis of *Wolbachia* status. For each subline, we measured maternal transmission by screening five female and five male F1s for *Wolbachia* using PCR. We extracted DNA from individual F1s using 96-well plates and a “squish” extraction buffer (10 mL Tris-HCl [1M], 0.0372 g EDTA, 0.1461 g NaCl, 90 mL dH20, followed by 150 mL Proteinase K after autoclaving) that allows for high-throughput and cost-effective DNA extraction. We screened each F1 for *Wolbachia* status using previously described PCR primers for the *Wolbachia* surface protein (*wsp*) and a second set of primers for the arthropod-specific 28S rDNA that served as a positive control^20,40,117^.

We computed maternal transmission for each genotype at each temperature as the mean proportion of *Wolbachia*-positive offspring produced by *Wolbachia*-positive mothers in each subline. We then estimated 95% bias-corrected and accelerated bootstrap (BC_a_) confidence intervals using the “boot.ci” function and 5000 iterations in the *boot* package in R^118,119^. We analyzed whether maternal transmission rates vary by *Wolbachia*-host genotype (e.g., *w*Mel-*D. melanogaster*) and temperature using a generalized linear mixed model (GLMM) logistic regression with the “glmer” function in the *lme4* R package^120^. We treated the *Wolbachia* status of F1s as the dependent variable and then included sex, the *Wolbachia*-host genotype, temperature, and a two-way interaction between the *Wolbachia*-host genotype and temperature as independent variables. We included sex in the model because we previously found that *w*Mel-like *Wolbachia* (*w*Mel and *w*Yak) are transmitted more efficiently to male offspring^20,21^. Finally, we included the subline of each F1 as a random effect. We assessed significance of the fixed effects with a likelihood ratio test (LRT) and type III sum of squares using the “mixed” function in the *afex* package^121^. Lastly, we used two-sided Wilcoxon rank-sum tests to evaluate whether transmission rates differ between 25° and 20°C for each strain.

### Wolbachia density in host ovaries

After each female was allowed to lay for eight days, we immediately dissected out her ovaries in chilled 1X PBS. Both the ovaries and remaining carcass were then frozen at -80°C. We then used qPCR to quantify *Wolbachia* density in the dissected ovaries and carcasses of each individual female. DNA was first extracted using a DNeasy Blood and Tissue Kit (Qiagen) and then amplified the *Wolbachia*-specific locus *ftsZ* and the *Drosophila*-specific locus *nAcRalpha-34E*. Preliminary analyses indicated that separate primers were needed for *Wolbachia* in the divergent A-Group (F: 5’-ATCCTTAACTGCGGCTCTTG-3’, R: 5’-TTCATCACAGCAGGAATGGG-3’) and B-Group lineages (F: 5’- CAGAGAAGCAAGAGCGGTAG-3’, R: 5’-TCTTCAAGTCCAAGCTCTGC -3’). We were able to use a single set of primers for all the *Drosophila* hosts (F: 5’- CTATGGTCGTTGACAGACT-3’, R: 5’-GTAGTACAGCTATTG TGGC-3’). We generated efficiency curves to confirm that each primer pair amplified with adequate efficiency for each *Wolbachia* strain and host species. The individual 10 μl reactions included 5 μl PowerUp SYBR Green Master Mix (Applied Biosystems), 0.25 μl of the forward and reverse primers, 0.5 μl of water, and 4 μl of gDNA. All qPCR reactions were amplified using the following cycling conditions: 50°C for 2 min, 95°C for 2 min, and then 40 cycles, with one cycle consisting of 95°C for 15 s, 58°C for 15 s, and 72°C for 1 min. We used the average cycle threshold (Ct) value of three technical replicates for each sample. We then estimated relative *Wolbachia* density as *2*^Δ^*^Ct^*, where *Δ*Ct* = *Ct*_nAcRalpha-34E_ − *Ct*_ftsZ_*^122^. We used a two-way ANOVA and type III sums of squares to test whether log-transformed *Wolbachia* density varied among *Wolbachia*-host genotypes and temperature for both the ovaries and the carcasses. We also used two-sided Wilcoxon rank-sum tests to test whether densities differed between 25° and 20°C for each tissue and *Wolbachia*-host genotype. If endoreplication in nurse cells is altered by temperature, changes to host ploidy in ovary tissues could plausibly influence our estimates of relative *Wolbachia* density (*2*^Δ*Ct*^) at each temperature^72^. We evaluated the raw Ct values for evidence of temperature differences in copy number at the host locus *nAcRalpha-34E* (Figure S6) and found that host ovary Ct values did not change significantly between temperature treatments, implying that differences in relative *Wolbachia* density are due to changes in absolute abundance of the *Wolbachia* gene.

### Cellular Wolbachia abundance in host oocytes

We used previously described protocols and confocal imaging to quantify cellular *Wolbachia* in host oocytes at each temperature^15,21^. Briefly, we paired 20 newly emerged virgin *Wolbachia*-positive females *en masse* with ten 3-5 day old virgin *Wolbachia*-negative males in individual vials for eight days to stimulate oocyte production. Females were then separated using CO_2_ and ovaries were dissected in a chilled dish of 1X PBS. Ovaries were fixed in 200 μl of devitellinizing solution (2% paraformaldehyde and 0.5% v/v NP40 in 1X PBS) mixed with 600 μl of heptane for 20 min on a shaker at room temperature. We then removed the organic layer using a brief centrifugation and washed the ovaries three times with PBS-T (0.1% Triton X-100 in 1X PBS), followed by three additional five min washes. Samples were treated with RNAse A (10 mg/ml) and left overnight at room temperature. The samples were then washed in PBS-T multiple times over two hours and then stained with a dilute solution of Alexa 488 phalloidin on a shaker for two hours for actin staining. Subsequently, the samples were washed again multiple times over the course of two more hour. Finally, the wash solution was removed and we added 60 μl of propidium iodide (PI) in mounting media (VECTASHIELD) and left again overnight. Ovaries were then mounted and carefully separated out again for ease of imaging. Slides were sealed with a coat of nail polish and stored at -20°C until imaging.

We used a Zeiss Olympia confocal microscope and the 40x Plan-Apochromat oil objective for image acquisition. Alexa 488 was imaged using the 488 nm laser line and emissions were collected from 493 to 584 nm. Propidium iodide was imaged using the 488 and 561 nm lines and emissions were collected from 584 to 718 nm. Stage 10 oocytes were identified based on two criterion. First, the egg chamber had to have two compartments, one with nurse cells and the other with the developing oocyte. Second, the oocyte had to account for approximately 50% of the egg chamber. The top and bottom of the oocyte were identified by scanning through the egg chamber and then a 0.8 μm slice was taken around the midplane. The pixel dwell time was 8.19 μsec.

Image analysis was conducted in the program Fiji^123^ following Russell *et al.*^63^. We first manually adjusted the contrast of each image to increase the threshold so that only white puncta corresponding to *Wolbachia* cells inside the oocyte were retained and all background noise was rendered black. We then used the polygon selection tool to select three different regions of the oocyte (see Figure S1 in Russell *et al.*): the whole oocyte, the posterior region, and the posterior cortex. We defined the posterior region as the posterior 1/8^th^ of the oocyte and the posterior cortex as the narrow cortical region where the pole plasm component Vasa is expressed^15,63^. The “area” and “integrated density” were measured for each region. Additionally, the average of a quadruplicate measure of “mean gray value” in background selections was measured to account for any remaining background fluorescence. The corrected total cell fluorescence (CTCF) was calculated as integrated density – (area x background mean gray value) for each region. We used two-way ANOVAs and type III sums of squares to test whether log-transformed CTCF values differed among *Wolbachia*-host genotypes and temperature in each region of the oocyte (whole oocyte, posterior region, and posterior cortex). We also used two-sided Wilcoxon rank-sum tests to test whether CTCF values differed between 25° and 20°C for each oocyte regions and *Wolbachia*-host genotype.

### Phylogenomic analyses

We conducted phylogenomic analyses to characterize the evolutionary relationships among *Wolbachia* strains and *Drosophila* host species. We obtained *Wolbachia* sequences from publicly available genome assemblies for *w*Mel^21^, *w*MelCS^66^, *w*Seg, *w*Cha^15^, *w*Ri^124^, *w*Ana^125^, *w*Aura, *w*Tri^51^, *w*Ha^126^, and *w*Mau^40^. We used Prokka 1.11 to identify homologs to known bacterial genes in the assemblies^127^. We avoided pseudogenes and paralogs in our phylogenetic analyses by selecting only single copy genes that uniquely matched a bacterial reference gene identified by Prokka. We removed loci with indels by requiring all homologs to have identical length in all *Wolbachia* genomes. A total of 170 full length single copy genes met these criteria, totaling 135,105 bp. We then estimated a Bayesian phylogram using RevBayes 1.0.8 under the GTR + Γ + I model partitioned by codon position^128^. Four independent runs were performed, which all converged on the same topology. All nodes were supported with Bayesian posterior probabilities >0.99.

To estimate an absolute chronogram for *Wolbachia*, we first estimated a relaxed-clock relative chronogram using RevBayes^128^ with the root age fixed to 1 using the GTR + Γ + I, partitioned by codon position, using the same birth-death prior as Turelli et al (2018)^51^. Each partition had an independent rate multiplier with prior Γ(1,1), as well as stationary frequencies and exchangeability rates drawn from flat, symmetrical Dirichlet distributions. For each branch, the branch-rate prior was Γ(7,7), normalized to a mean of 1 across all branches. We performed four independent runs that all agreed. We converted the relative chronogram into an absolute chronogram using the scaled distribution Γ(7,7) × 6.87 × 10^−9^ substitutions per third position site per year, derived from the posterior distribution estimated by Richardson et al. (2012)^85^, assuming 10 generations per year. Absolute branch lengths were calculated as the relative branch length times the third position rate multiplier divided by the substitutions per third-position site per year estimate above.

We used similar methods to generate a phylogram for the *Drosophila* host species. Publicly available sequences were obtained for *D. melanogaster*^129^, *D. simulans*^130^, *D. mauritiana*^40^, *D. seguyi*, *D. chauvacae*^131^, *D. auraria*, *D. triauraria*^51^, and *D. ananassae*^132^. A host phylogeny was generated using the same nuclear genes implemented in Turelli *et al.*^51^: *aconitase, aldolase, bicoid, ebony, enolase, esc, g6pdh, glyp, glys, ninaE, pepck, pgi, pgm, pic, ptc, tpi, transaldolase, white, wingless,* and *yellow*. We used BLAST with the *D. melanogaster* coding sequences to extract orthologs from the genomes of each host species. Sequences were then aligned with MAFFT 7^133^. Finally, we used RevBayes and the GTR + Γ + I model partitioned by codon position and gene to accommodate potential variation in the substitution process among genes. All nodes were supported with Bayesian posterior probabilities of 1.

The resulting phylograms were used to test whether *Wolbachia* density and oocyte cellular abundance exhibit phylogenetic signal on either the *Wolbachia* or host phylogenies. For the host phylogeny, we pooled estimates of *Wolbachia* density and oocyte abundance by host species, because two of the hosts (*D. melanogaster* and *D. simulans*) had data for multiple genotypes carrying different *Wolbachia* strains. We used our estimates of *Wolbachia* density (2*^ΔCt^*) and cellular oocyte abundance (log-transformed CTCF) to test for phylogenetic signal using Pagel’s lambda (λ)^68^. A Pagel’s λ of 0 indicates that character evolution occurs independently of phylogenetic relationships, whereas λ = 1 is consistent with a Brownian motion model of character evolution. We used the “phylosig” function and a likelihood ratio test (LRT) in the *phytools* package^134^ to compare the fitted value to a model assuming no phylogenetic signal (λ = 0). We also used a Monte Carlo-based method to generate 95% confidence intervals surrounding our estimates using 1,000 bootstrap replicates in the *pmc* package^135^.

### Statistics and reproducibility

All statistical analyses were performed in R^119^. Specific details for each statistical analysis are reported above. We conducted distribution and leverage analyses to evaluate assumptions of normality and then used data transformations and non-parametric tests when applicable (described above). Samples sizes and test statistics for each *Wolbachia*-host genotype are reported in the Supplementary Information. For each experiment, we collected the largest sample size that was feasible within our experimental design. We do not report any issues with reproducibility.

## ACKNOWLEDGEMENTS

We are especially thankful to W. Sullivan for very useful discussions that improved the quality of this manuscript. We thank W. Conner for assistance with bioinformatic analyses. J. Statz provided thoughtful feedback on the manuscript. Support and instrumentation was provided by the Cellular Genetics Ecosystem in the Genomics Core, the Environmental Control for Organismal Research (ECOR) facility, and the Center for Biomolecular Structure and Dynamics at the University of Montana. This study was funded by a National Science Foundation CAREER (2145195) Award to BSC. BSC was also supported by a National Institute of General Medical Sciences of the National Institutes of Health MIRA Award (R35GM124701).

## AUTHOR CONTRIBUTIONS

M.T.J.H. designed the project, collected data, performed analyses, and prepared the manuscript. T.B.W. collected data. B.S.C. acquired funding, designed the project, prepared the manuscript, and provided leadership. All authors reviewed and edited the manuscript prior to submission.

## COMEPETING INTERESTS

The authors declare no competing interests.

## DATA AVAILABILITY

All source data on the estimation of maternal transmission, *Wolbachia* densities, and cellular *Wolbachia* abundance are included as Supplementary Data files.

