## Supplemental Information for "Comparative analysis of *Wolbachia* maternal transmission and localization in host ovaries"

### SUPPLEMENTARY INFORMATION

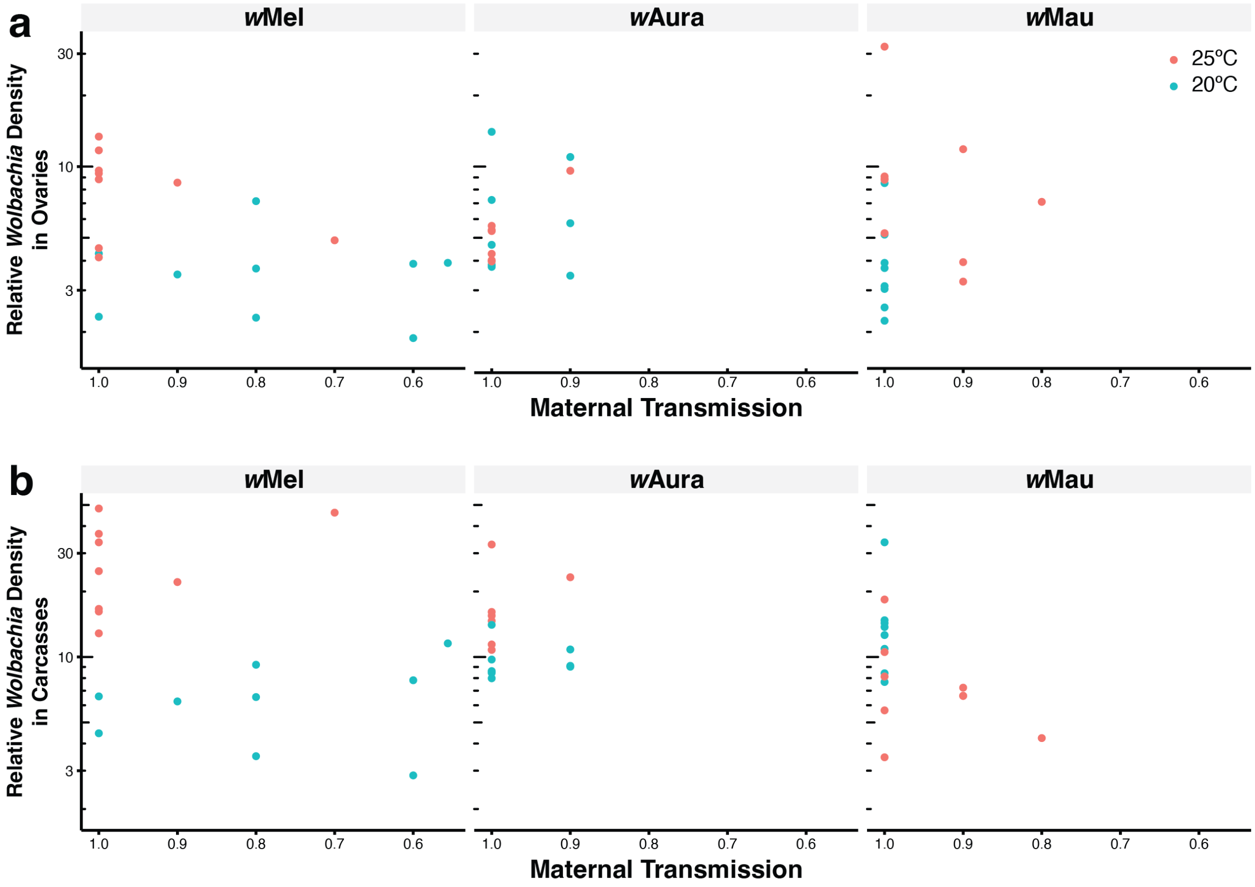


**Figure S1.** Correlation plots showing the relationship between maternal transmission rates and *Wolbachia* density in the **(a)** ovaries and **(b)** carcasses of the same individual females. Plots are shown for the three *Wolbachia* strains that exhibited significant variation in transmission by their host species across temperatures: *w*Mel, *w*Aura, and *w*Mau

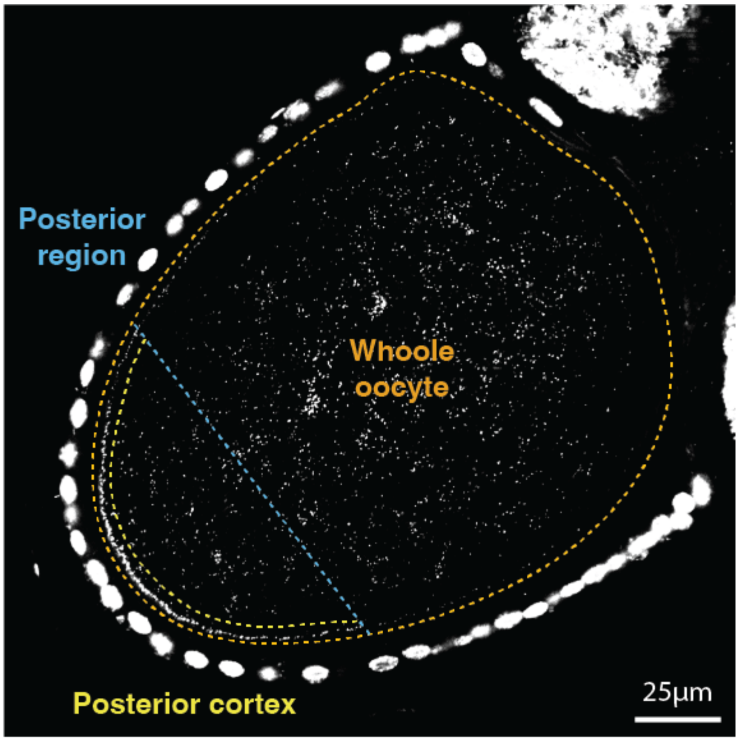


**Figure S2.** Quantification of cellular *Wolbachia* abundance in stage 10 oocytes measured as *Wolbachia* fluorescence due to propidium iodide (CTCF) using the PI greyscale channel. Schematic shows each region of the oocyte: the whole oocyte, the posterior region, and the posterior cortex.

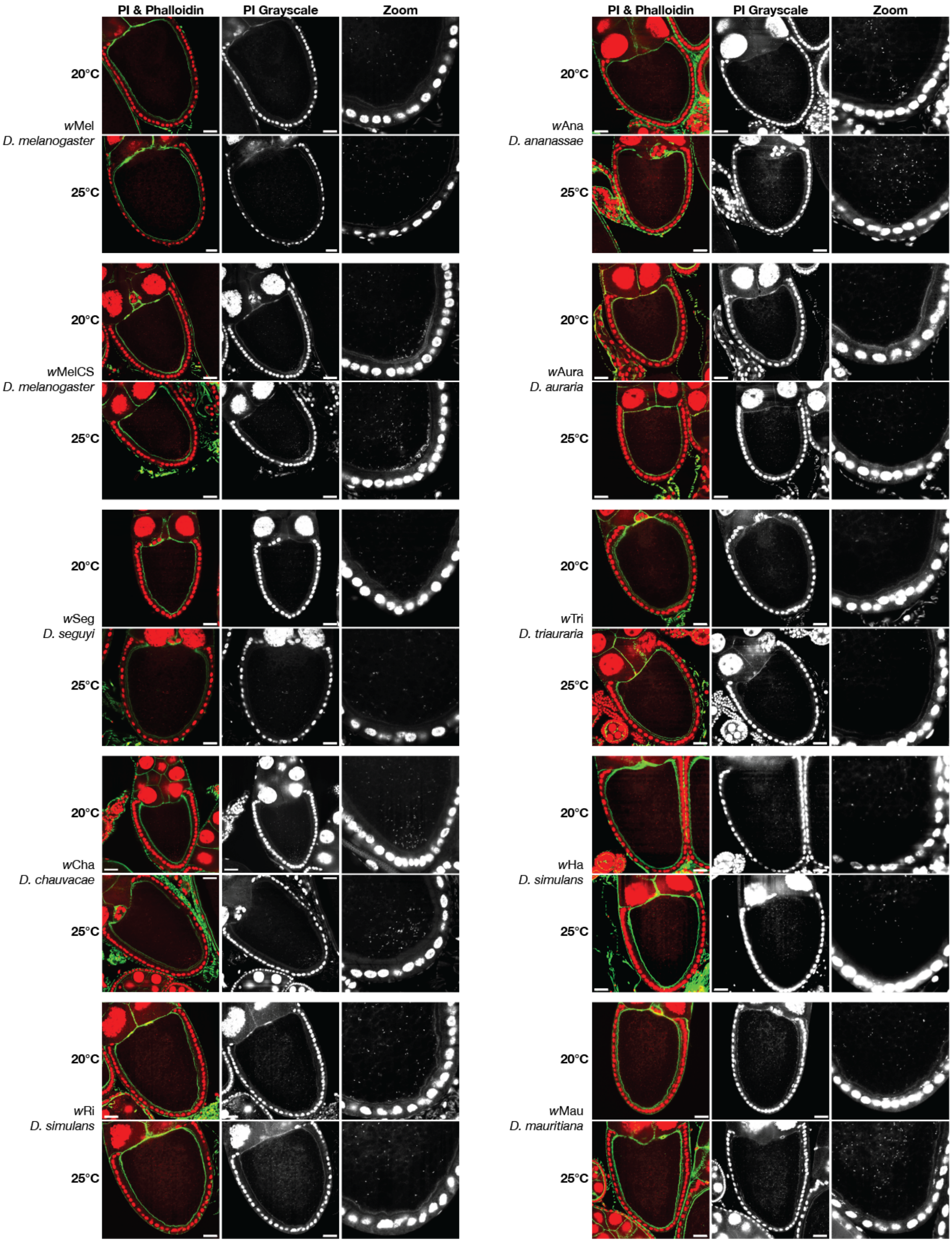


**Figure S3.** Representative confocal images of stage 10 oocytes for each *Wolbachia*-host genotype at 25° and 20°C. Samples are stained with propidium iodide (PI) for DNA (red) and phalloidin for actin (green). Grayscale images of the PI channel are shown in the middle column and enlarged images of the oocyte posterior region are shown in the third column for each genotype. Scale bars are set to 25 µm.

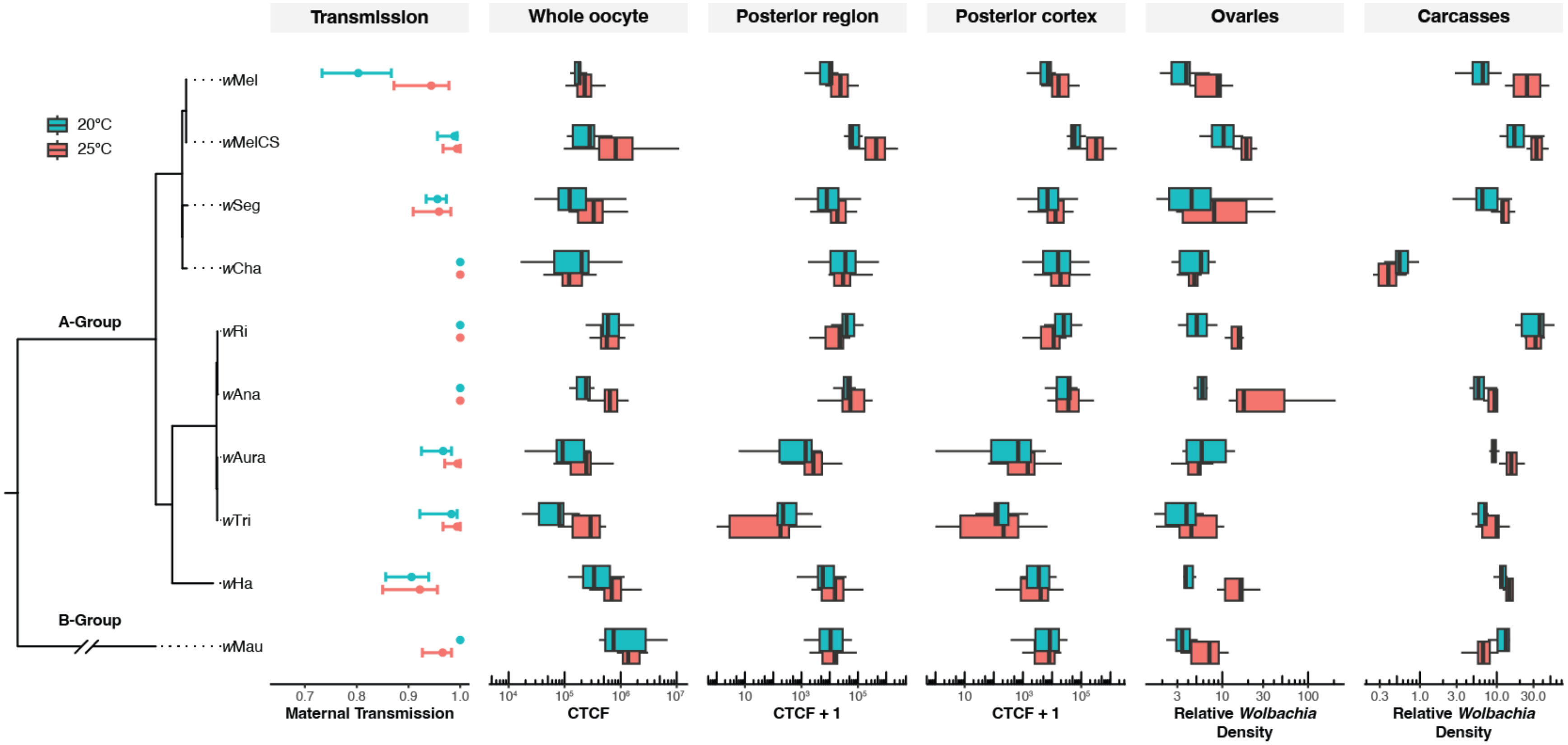


**Figure S4.** Summary of all data in relation to the *Wolbachia* phylogeny.

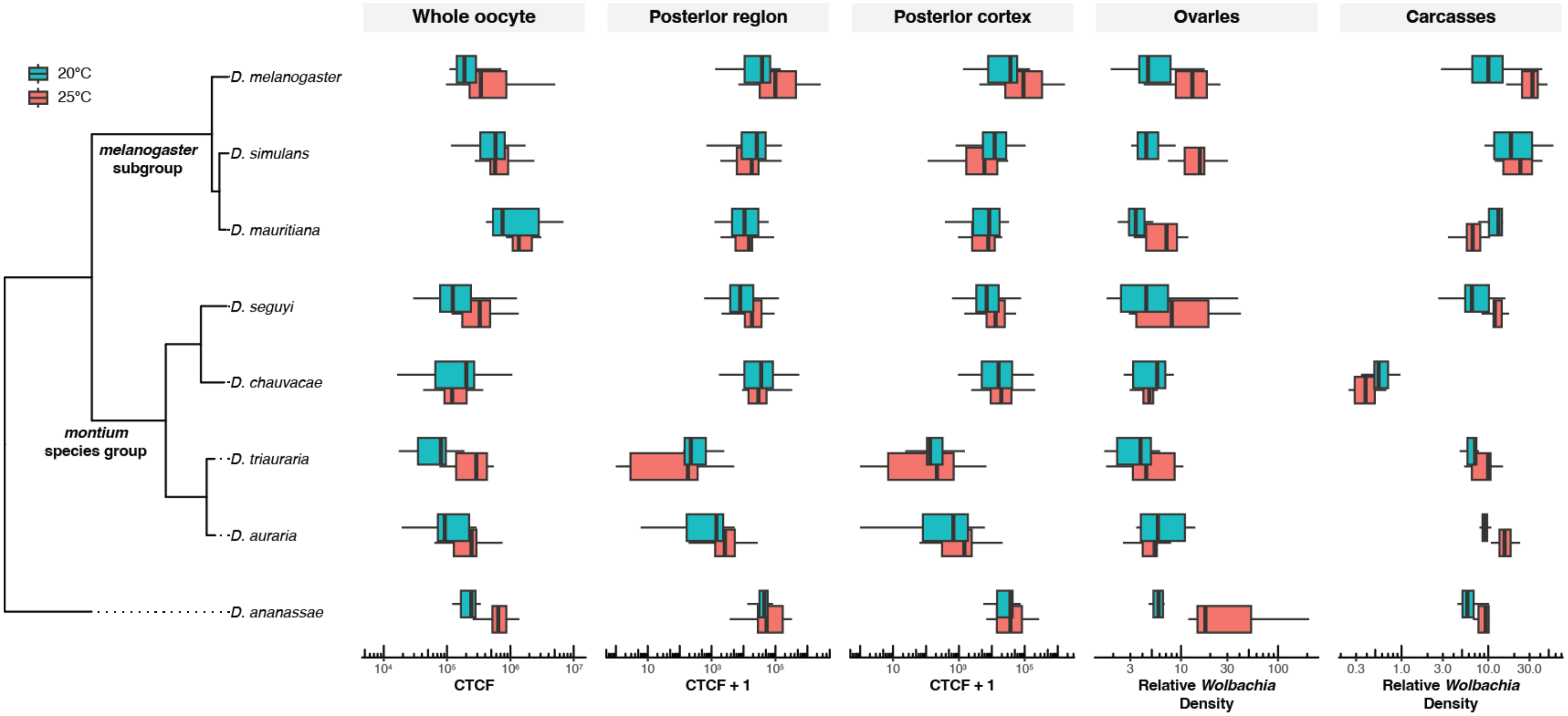


**Figure S5.** Summary of all data in relation to the *Drosophila* host phylogeny.

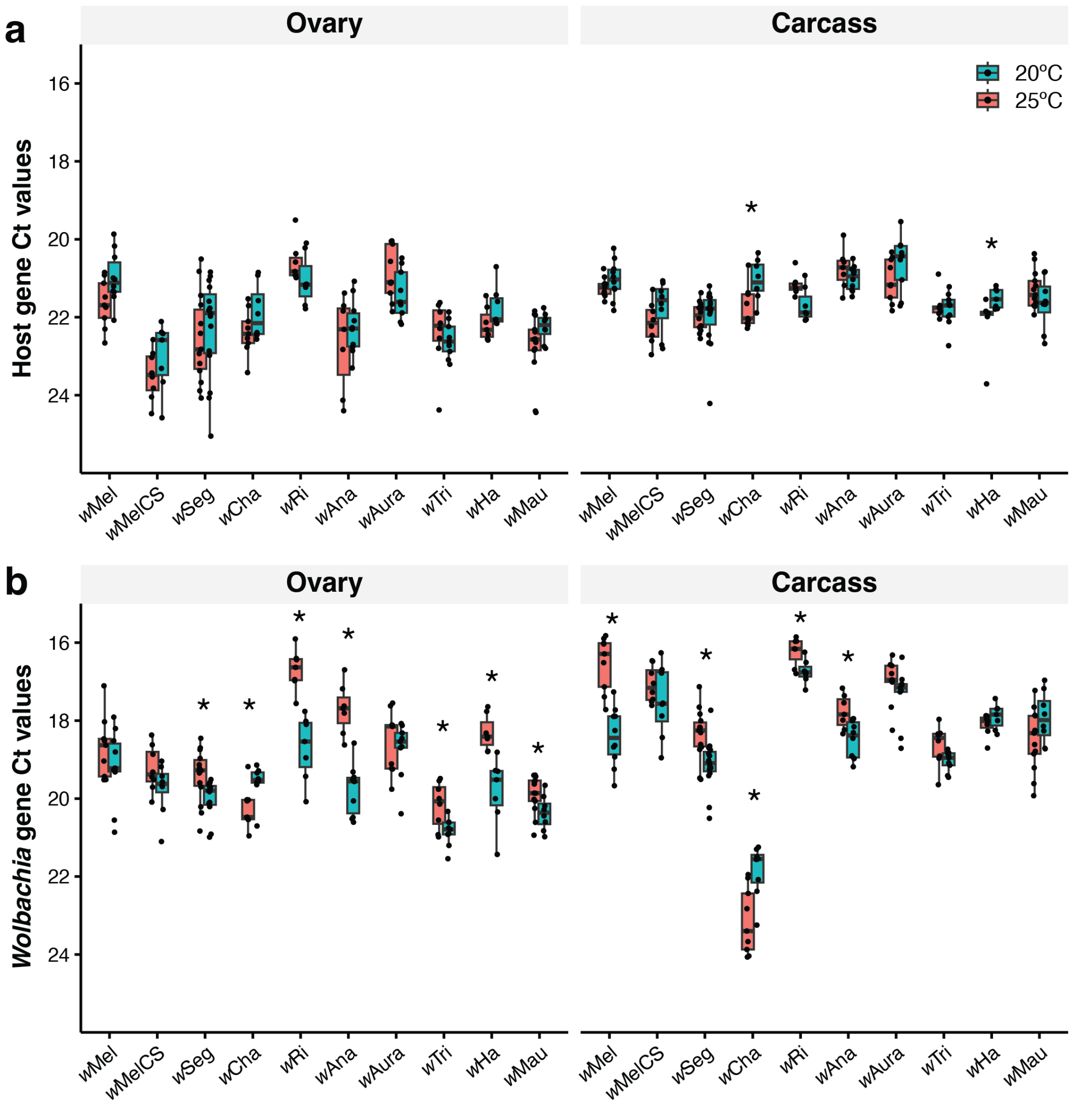


**Figure S6.** Raw Ct values from qPCR for **(a)** the host gene *nAcRalpha-34E* and **(b)** the *Wolbachia* gene *ftsZ*. Note that Ct values on the y-axes are reversed, because lower Ct values correspond to higher abundance of the gene. Asterisks indicate that the Ct values differ between 25° and 20°C for a given *Wolbachia*-host genotype according to a Wilcoxon rank-sum test at $P < 0.05$.

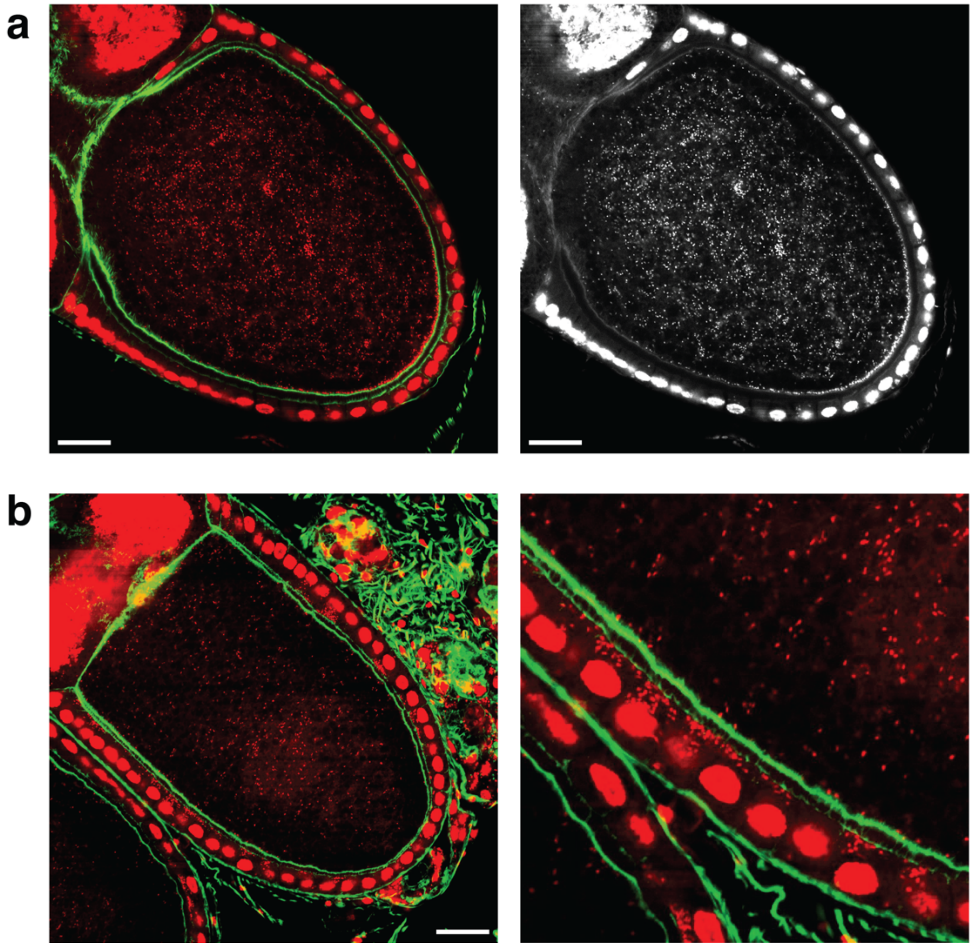


**Figure S7. (a)** Confocal image of a *D. melanogaster* stage 10 oocyte with a particularly high cellular abundance of *w*MelCS. DNA (PI) is shown in red and actin (phalloidin) is shown in green. To the right, a single channel image of the PI stain is shown. **(b)** Confocal image of a *D. mauritiana* stage 10 oocyte with many *w*Mau cells in the follicle cells. An enlarged image of the follicle cells is shown on the right. Scale bars are set to 25 µm.

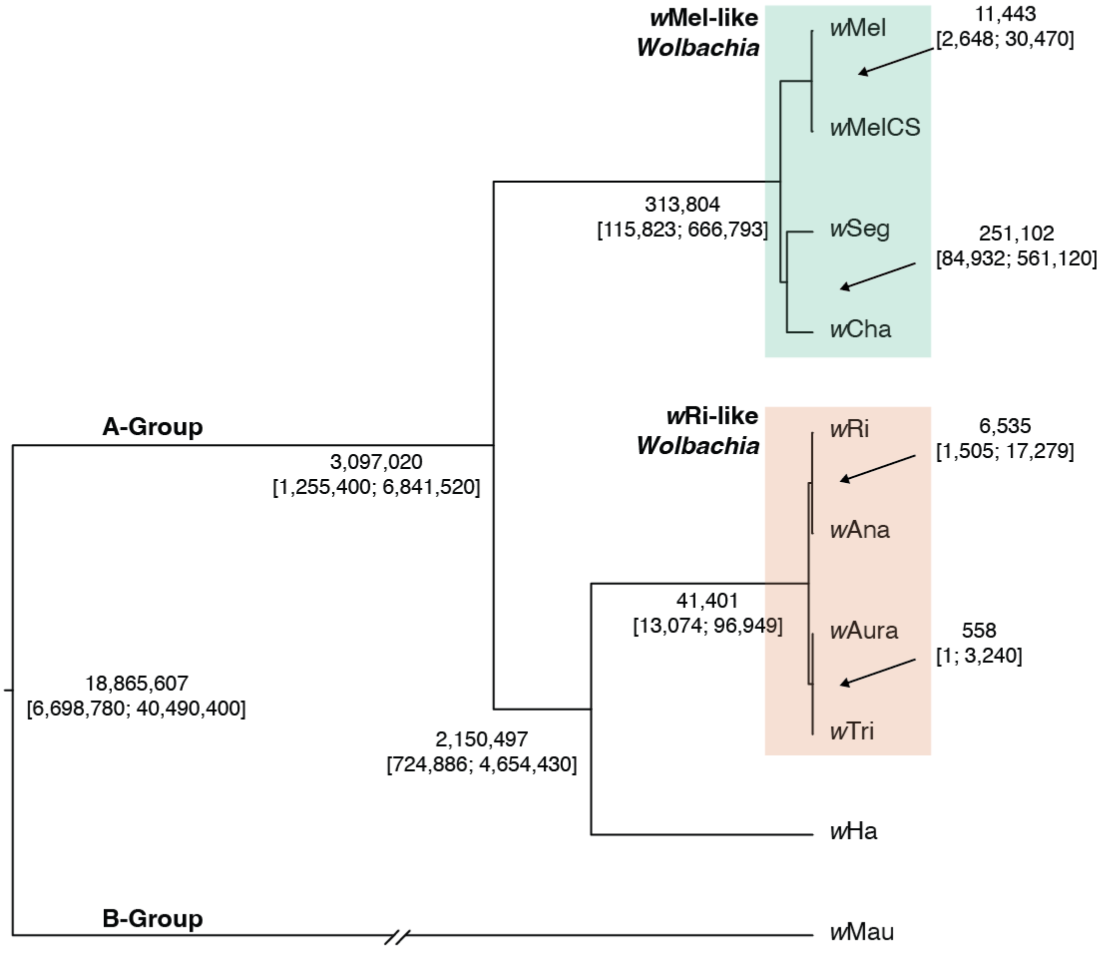


**Figure S8.** Bayesian chronogram with absolute age estimates (and 95% credible intervals) based on a calibration of rates of *Wolbachia* divergence from Richardson et al., (2012)[1]. The chronogram was estimated using 170 full length single copy genes. All nodes are supported with Bayesian posterior probabilities of 1.

**Table S1.** Summary of fly lines used in maternal transmission experiments that crossed *Wolbachia*-positive females with *Wolbachia*-free males.

| ***Wolbachia*** | **Species** | ***Wolbachia*-positive Genotype** | ***Wolbachia*-free genotype (TC = tetracycline-cleared)** |
|---|---|---|---|
| *w*Mel | *D. melanogaster* | melHB22 | mel17IN10B7 |
| *w*MelCS | *D. melanogaster* | melCSBerk | mel17IN10B7 |
| *w*Seg | *D. seguyi* | segL1 | segL2-TC |
| *w*Cha | *D. chauvacae* | chauvL1 | chauvL2-TC |
| *w*Ri | *D. simulans* | simR84 | sim299 |
| *w*Ana | *D. ananassae* | anaRC102 | ana14024 |
| *w*Aura | *D. auraria* | auraL5 | auraL2-TC |
| *w*Tri | *D. triauraria* | triL1 | triE-15301 |
| *w*Ha | *D. simulans* | simCar5 | sim299 |
| *w*Mau | *D. mauritiana* | mauR21 | mauR4 |

**Table S2.** Imperfect maternal transmission ($\mu$) is estimated as the proportion of *Wolbachia*-free offspring produced by *Wolbachia*-positive females. Mean estimates of imperfect maternal transmission (± $BC_a$ confidence intervals) are shown for 25° and 20°C. The number of sublines (*N*) used to generate means are shown for each genotype and temperature.

| ***Wolbachia*** | **Species** | **Treatment** | ***N* sublines** | ***μ*** | **Female *μ*** | **Male *μ*** |
|---|---|---|---|---|---|---|
| *w*Mel | *D. melanogaster* | 20°C | 18 | 0.197 [0.133, 0.269] | 0.322 [0.2, 0.493] | 0.078 [0.044, 0.133] |
| *w*Mel | *D. melanogaster* | 25°C | 18 | 0.056 [0.022, 0.128] | 0.096 [0.033, 0.237] | 0.011 [0, 0.067] |
| *w*MelCS | *D. melanogaster* | 20°C | 18 | 0.011 [0.006, 0.05] | 0 [NA, NA] | 0.022 [0.011, 0.089] |
| *w*MelCS | *D. melanogaster* | 25°C | 18 | 0.006 [0, 0.039] | 0.011 [0, 0.067] | 0 |
| *w*Seg | *D. seguyi* | 20°C | 23 | 0.044 [0.026, 0.068] | 0.072 [0.037, 0.12] | 0.017 [0.009, 0.078] |
| *w*Seg | *D. seguyi* | 25°C | 22 | 0.041 [0.018, 0.095] | 0.082 [0.036, 0.182] | 0 |
| *w*Cha | *D. chauvacae* | 20°C | 17 | 0 | 0 | 0 |
| *w*Cha | *D. chauvacae* | 25°C | 18 | 0 | 0 | 0 |
| *w*Ri | *D. simulans* | 20°C | 19 | 0 | 0 | 0 |
| *w*Ri | *D. simulans* | 25°C | 18 | 0 | 0 | 0 |
| *w*Ana | *D. ananassae* | 20°C | 18 | 0 | 0 | 0 |
| *w*Ana | *D. ananassae* | 25°C | 18 | 0 | 0 | 0 |
| *w*Aura | *D. auraria* | 20°C | 12 | 0.033 [0.017, 0.075] | 0.045 [0.017, 0.112] | 0.017 [0, 0.1] |
| *w*Aura | *D. auraria* | 25°C | 20 | 0.005 [0, 0.035] | 0 | 0.008 [0, 0.05] |
| *w*Tri | *D. triauraria* | 20°C | 18 | 0.017 [0.006, 0.061] | 0.033 [0.011, 0.144] | 0 |
| *w*Tri | *D. triauraria* | 25°C | 18 | 0.006 [0, 0.039] | 0.011 [0, 0.067] | 0 |
| *w*Ha | *D. simulans* | 20°C | 18 | 0.094 [0.067, 0.144] | 0.158 [0.1, 0.233] | 0.033 [0.011, 0.149] |
| *w*Ha | *D. simulans* | 25°C | 18 | 0.078 [0.044, 0.156] | 0.144 [0.078, 0.3] | 0.011 [0, 0.056] |
| *w*Mau | *D. mauritiana* | 20°C | 18 | 0 | 0 | 0 |
| *w*Mau | *D. mauritiana* | 25°C | 18 | 0.034 [0.017, 0.073] | 0.069 [0.033, 0.147] | 0 |

**Table S3.** Wilcoxon rank-sum tests testing for a difference in maternal transmission between 25° and 20°C for each *Wolbachia*-host genotype.

| ***Wolbachia*** | **Species** | ***W*** | ***P*** |
|---|---|---|---|
| *w*Mel | *D. melanogaster* | 251.5 | 0.003 |
| *w*MelCS | *D. melanogaster* | 171 | 0.574 |
| *w*Seg | *D. seguyi* | 286 | 0.381 |
| *w*Cha | *D. chauvacae* | NA | NA |
| *w*Ri | *D. simulans* | NA | NA |
| *w*Ana | *D. ananassae* | NA | NA |
| *w*Aura | *D. auraria* | 154 | 0.038 |
| *w*Tri | *D. triauraria* | 171.5 | 0.553 |
| *w*Ha | *D. simulans* | 188 | 0.386 |
| *w*Mau | *D. mauritiana* | 117 | 0.019 |

**Table S4.** Wilcoxon rank-sum tests testing for a differences in *Wolbachia* density between 25° and 20°C in female ovaries and carcasses for each *Wolbachia*-host genotype. The number of females (*N*) included in each analysis is shown.

| | | | **Ovaries** | | **Carcasses** | |
|---|---|---|---|---|---|---|
| ***Wolbachia*** | **Species** | ***N*** | ***W*** | ***P*** | ***W*** | ***P*** |
| *w*Mel | *D. melanogaster* | 19 | 84 | 0.001 | 90 | <0.001 |
| *w*MelCS | *D. melanogaster* | 16 | 45 | 0.054 | 53 | 0.028 |
| *w*Seg | *D. seguyi* | 29 | 145 | 0.085 | 171 | 0.003 |
| *w*Cha | *D. chauvacae* | 17 | 26 | 0.370 | 15 | 0.046 |
| *w*Ri | *D. simulans* | 14 | 48 | 0.001 | 26 | 0.902 |
| *w*Ana | *D. ananassae* | 17 | 70 | <0.001 | 54 | 0.070 |
| *w*Aura | *D. auraria* | 18 | 33 | 0.546 | 77 | <0.001 |
| *w*Tri | *D. triauraria* | 17 | 46 | 0.370 | 43 | 0.252 |
| *w*Ha | *D. simulans* | 14 | 49 | 0.001 | 40 | 0.053 |
| *w*Mau | *D. mauritiana* | 22 | 89 | 0.006 | 12 | 0.003 |

**Table S5.** Wilcoxon rank-sum tests testing for a differences in cellular *Wolbachia* abundance between 25° and 20°C in stage 10 oocytes for each *Wolbachia*-host genotype. Tests are shown for the whole oocyte, the posterior region, and the posterior cortex. The number of oocytes (*N*) included in each analysis is shown.

| | | | **Whole oocyte** | | **Posterior region** | | **Posterior cortex** | |
|---|---|---|---|---|---|---|---|---|
| ***Wolbachia*** | **Species** | ***N*** | ***W*** | ***P*** | ***W*** | ***P*** | ***W*** | ***P*** |
| *w*Mel | *D. melanogaster* | 24 | 99 | 0.119 | 113 | 0.015 | 122 | 0.002 |
| *w*MelCS | *D. melanogaster* | 30 | 189 | 0.001 | 203 | <0.001 | 205 | <0.001 |
| *w*Seg | *D. seguyi* | 35 | 216 | 0.020 | 198 | 0.089 | 197 | 0.096 |
| *w*Cha | *D. chauvacae* | 30 | 112 | 1 | 113 | 1 | 130 | 0.4864 |
| *w*Ri | *D. simulans* | 30 | 101 | 0.653 | 45 | 0.004 | 49 | 0.008 |
| *w*Ana | *D. ananassae* | 30 | 218 | <0.001 | 142 | 0.233 | 127 | 0.567 |
| *w*Aura | *D. auraria* | 31 | 176 | 0.027 | 172 | 0.041 | 150 | 0.244 |
| *w*Tri | *D. triauraria* | 29 | 187 | <0.001 | 79 | 0.265 | 103.5 | 0.965 |
| *w*Ha | *D. simulans* | 22 | 92 | 0.036 | 80 | 0.203 | 53 | 0.674 |
| *w*Mau | *D. mauritiana* | 22 | 81 | 0.193 | 66 | 0.748 | 53 | 0.652 |

**Table S6**. Tests for phylogenetic signal using Pagel's λ. Estimates of λ (± 95% confidence intervals) are shown with *P* values from LRTs for *Wolbachia* density in host tissues (top) and cellular abundance in host oocytes (bottom) at each temperature. λ values were estimated using both the *Wolbachia* and host phylograms (Figure 1).

| | | **Ovary *Wolbachia* density** | | | | **Carcasse *Wolbachia* density** | | | |
|---|---|---|---|---|---|---|---|---|---|
| **Phylogeny** | **Temperature** | **λ** | **Lower CI** | **Upper CI** | ***P*** | **λ** | **Lower CI** | **Upper CI** | ***P*** |
| *Wolbachia* | 25°C | 0.000 | 0.000 | 0.670 | 1.000 | 0.000 | 0.000 | 0.617 | 1.000 |
| | 20°C | 0.000 | 0.000 | 0.674 | 1.000 | 0.000 | 0.000 | 0.622 | 1.000 |
| Host | 25°C | 1.000 | 0.000 | 1.000 | 0.001 | 0.000 | 0.000 | 0.920 | 1.000 |
| | 20°C | 0.000 | 0.000 | 0.833 | 1.000 | 0.616 | 0.000 | 1.000 | 0.288 |

| | | **Whole oocyte *Wolbachia* abundance** | | | | **Posterior region *Wolbachia* abundance** | | | | **Posterior cortex *Wolbachia* abundance** | | | |
|---|---|---|---|---|---|---|---|---|---|---|---|---|---|
| **Phylogeny** | **Temperature** | **λ** | **Lower CI** | **Upper CI** | ***P*** | **λ** | **Lower CI** | **Upper CI** | ***P*** | **λ** | **Lower CI** | **Upper CI** | ***P*** |
| *Wolbachia* | 25°C | 0.000 | 0.000 | 0.702 | 1.000 | 0.000 | 0.000 | 0.694 | 1.000 | 0.000 | 0.000 | 0.602 | 1.000 |
| | 20°C | 0.472 | 0.000 | 0.910 | 0.521 | 0.000 | 0.000 | 0.72046 | 1.000 | 0.000 | 0.000 | 0.674 | 1.000 |
| Host | 25°C | 0.736 | 0.000 | 1.000 | 0.119 | 0.000 | 0.000 | 0.819 | 1.000 | 0.000 | 0.000 | 0.861 | 1.000 |
| | 20°C | 0.644 | 0.000 | 1.000 | 0.250 | 1.000 | 0.000 | 1.000 | 0.037 | 1.000 | 0.000 | 1.000 | 0.036 |
